# Molecular subtypes of high-grade serous ovarian cancer across racial groups and gene expression platforms

**DOI:** 10.1101/2023.11.01.565179

**Authors:** Natalie R. Davidson, Mollie E. Barnard, Ariel A. Hippen, Amy Campbell, Courtney E. Johnson, Gregory P. Way, Brian K. Dalley, Andrew Berchuck, Lucas A. Salas, Lauren C. Peres, Jeffrey R. Marks, Joellen M. Schildkraut, Casey S. Greene, Jennifer A. Doherty

**Author notes:** Corresponding authors: 1. Mollie E. Barnard, ScD 72 E. Concord St. Boston, MA, 02118, 2. Natalie R. Davidson, PhD, Department of Biomedical Informatics University of Colorado, Anschutz Medical Campus 1890 N Revere Ct, Aurora, CO. **Conflict of interest declaration:** LCP received grant funding from Bristol Myers Squibb for work external to this manuscript. The remaining authors declare no potential conflicts of interest. Denotes equal contributions (Natalie R. Davidson, Mollie E. Barnard). Denotes equal contributions (Joellen M. Schildkraut, Casey S. Greene, Jennifer A. Doherty).

## Abstract

**Introduction:** High-grade serous carcinoma (HGSC) gene expression subtypes are associated with differential survival. We characterized HGSC gene expression in Black individuals and considered whether gene expression differences by race may contribute to poorer HGSC survival among Black versus non-Hispanic White individuals.

**Methods:** We included newly generated RNA-Seq data from Black and White individuals, and array-based genotyping data from four existing studies of White and Japanese individuals. We assigned subtypes using K-means clustering. Cluster- and dataset-specific gene expression patterns were summarized by moderated t-scores. We compared cluster-specific gene expression patterns across datasets by calculating the correlation between the summarized vectors of moderated t-scores. Following mapping to The Cancer Genome Atlas (TCGA)-derived HGSC subtypes, we used Cox proportional hazards models to estimate subtype-specific survival by dataset.

**Results:** Cluster-specific gene expression was similar across gene expression platforms. Comparing the Black study population to the White and Japanese study populations, the immunoreactive subtype was more common (39% versus 23%-28%) and the differentiated subtype less common (7% versus 22%-31%). Patterns of subtype-specific survival were similar between the Black and White populations with RNA-Seq data; compared to mesenchymal cases, the risk of death was similar for proliferative and differentiated cases and suggestively lower for immunoreactive cases (Black population HR=0.79 [0.55, 1.13], White population HR=0.86 [0.62, 1.19]).

**Conclusions:** A single, platform-agnostic pipeline can be used to assign HGSC gene expression subtypes. While the observed prevalence of HGSC subtypes varied by race, subtype-specific survival was similar.

**Statement of Significance:** A single pipeline was used to subtype ovarian high-grade serous carcinoma (HGSC) with array-based or RNA-Seq gene expression data. Subtype distributions differed by race, but subtype-specific survival was similar across racial groups.

## Introduction

Ovarian cancer is a highly fatal malignancy comprised of multiple histologically-defined subtypes (i.e., “histotypes”). High-grade serous carcinoma (HGSC) is the most common histotype (1), and an important contributor to ovarian cancer mortality (2–4). HGSC is also molecularly heterogeneous, which is relevant to both prognosis and treatment. Prior studies have described between three and five molecularly distinct subtypes (5–10), and, while no gold standard exists for defining these subtypes, they are commonly mapped to the four TCGA-derived subtypes, which are similar to those reported in Tothill et al., 2008: mesenchymal (Tothill C1.MES), proliferative (Tothill C5.PRO), immunoreactive (Tothill C2.IMM), and differentiated (Tothill C4.DIF) (5,11). Key characteristics of the mesenchymal subtype include high expression of HOX genes, increased stromal components, and poor survival (5,9,11). The proliferative subtype has been characterized by low expression of ovarian tumor markers (e.g., MUC1, MUC16), high expression of transcription factors, and intermediate survival (5,9,11).

Defining characteristics of the immunoreactive subtype include the enrichment of genes and pathways associated with an immune response, including CD3+/CD8+ T-cell markers and genes in the CXCL9, CXCL10, CXCL11/CXCR3 axis for immune activation, and more favorable survival (5,9,11). The differentiated subtype has been the most difficult to characterize and reproduce (7,12), yet multiple studies have described it as having high expression of ovarian tumor markers (e.g., MUC1, MUC16, SLPI) and intermediate to good survival (5,9).

Since HGSC gene expression-based subtypes have potential clinical utility, it is of interest to develop a clinical-grade, gene expression-based subtype classifier that is applicable across diverse populations. One recently developed gene-set assay, the Predictor of high-grade-serous Ovarian carcinoma molecular subTYPE (PrOTYPE) assay, was derived using array-based gene expression data from mostly non-Hispanic White individuals and migrated to a NanoString platform (11). The PrOTYPE assay has been reported to classify HGSC into the four TCGA-derived subtypes with >95% accuracy (11). Here, we leverage RNA-sequencing (RNA-seq) data to characterize HGSC gene expression subtypes among a more racially diverse study population, including a population of >300 self-identified Black individuals. Prior studies have noted that Black individuals have poorer overall ovarian cancer survival (13), and poorer three-year and six-year HGSC survival when compared to White individuals (14,15), so inclusion of Black individuals in the derivation of HGSC molecular subtypes is especially important.

In the present study, we used K-means clustering to assign HGSC tumors from Black, White, and Japanese individuals to molecular subtypes (7). Following subtype assignment, we compared subtype-specific gene expression, subtype frequency, and subtype-specific survival across all possible pairs of studies with different racial distributions (e.g., Black, White, Japanese), and different data types (array-based gene expression data, and RNA-seq).

## Methods

### Primary study population

We included epithelial ovarian cancer cases enrolled in one of two population-based case-control studies, the North Carolina Ovarian Cancer Study (NCOCS, diagnosis dates 1999-2005) (16), and the African American Cancer Epidemiology Study (AACES, diagnosis dates 2010-2015) (15,17). Both studies enrolled epithelial ovarian cancer cases covering a range of histotypes, grades, and stages, though some of the most aggressive cases were missed because they were feeling very ill or were already deceased by the time they were invited to participate in research (15). Written informed consent was obtained for NCOCS participants, while AACES participants provided verbal consent and signed medical record and pathology release forms to allow for access to tumor tissue. All cases were confirmed via centralized pathology review. Both the NCOCS and AACES studies were approved by the Duke Medical Center Institutional Review Board (IRB) and the IRBs of participating enrollment sites.

Together, the AACES and NCOCS included 747 Black ovarian cancer cases, 464 of which were HGSC. Of these, 325 provided consent to participate in biospecimen-based research and had adequate tissue available to pursue RNA extraction (**Supplemental Figure 1A**). Fifty-three of these cases were excluded from gene expression analyses due to a history of neoadjuvant chemotherapy, which can influence observed gene expression (**Supplemental Figure 1B**).

Following these exclusions, there were 272 self-identified Black or African American cases (3 Hispanic; 269 non-Hispanic) who we subsequently refer to as “SchildkrautB”. The NCOCS study included 1,014 White cases, 484 of which were HGSC. Of these, 316 provided consent to participate in biospecimen-based research and had sufficient tissue available to pursue RNA extraction (**Supplemental Figure 2A**). None of these cases had neoadjuvant chemotherapy prior to tissue collection, so all were considered eligible for gene expression analyses. We subsequently refer to this set of 316 non-Hispanic White cases as “SchildkrautW”.

Demographic characteristics, disease characteristics, and vital status were available for all individuals included in SchildkrautB and SchildkrautW. Information on age at diagnosis was obtained from questionnaires and pathology reports. Tumor stage, debulking status, and use of neoadjuvant chemotherapy were abstracted from medical records and pathology reports. The proportions of intratumoral CD3+ T cells and CD3+/CD8+ suppressor T cells were obtained from multiplex immunofluorescence staining of formalin-fixed paraffin-embedded (FFPE) tissue (18). Vital status was assessed using data from population-based cancer registries, obituaries, LexisNexis, and the National Death Index.

### Acquisition of gene expression data

RNA was extracted from FFPE tumor tissue and stored at −80°C. An initial quality control (QC) evaluation revealed substantial RNA degradation, so a re-purification step consisting of DNAase treatment and purification on a Zymo research spin column was completed before library preparation to reduce the bulk of degraded RNA product (i.e., RNA product <200 nucleotides in length). Following re-purification, RNA libraries were prepared from total RNA samples (5-100 ng) using reagents from the Illumina Stranded mRNA Prep (cat# 20020189) and the Illumina RNA UD Indexes Set (20091657) for reverse transcription, adapter ligation, and PCR amplification. Amplified libraries were hybridized to biotin-labeled probes from the Illumina Exome Panel (cat# 20020183) using the Illumina RNA Fast Hyb Enrichment kit (20040540) to generate strand-specific libraries enriched for coding regions of the transcriptome. The quality of exon-enriched libraries was assessed on an Agilent Technologies 2200 TapeStation using a D1000 ScreenTape assay (cat# 5067-5582 and 5067-5583). The molarity of adapter-modified molecules was defined by quantitative PCR using the Kapa Biosystems Kapa Library Quant Kit (cat#KK4824). Individual libraries were normalized to 0.95 nM in preparation for Illumina sequence analysis. Sequencing libraries were chemically denatured and applied to an Illumina NovaSeq flow cell using the NovaSeq XP workflow (20043131). Following the transfer of the flow cell to an Illumina NovaSeq 6000 instrument, a 150 x 150 cycle paired-end sequence run was performed using a NovaSeq 6000 S4 reagent Kit v1.5 (20028312).

### Quantification of gene expression data

We trimmed adapters and filtered read quality using fastp (19). We filtered to reads with a PHRED score of at least 15 and a length of at least 20 base pairs. While a minimum PHRED score of 15 may include some reads with low quality, we found that most bases across all samples in SchildkrautB and SchildkrautW had a quality score greater than 30 (**Supplemental Figure 3**). We quantified paired-end reads with Salmon (version 1.4.0) (20) using GRCh38 release 95. We used the seqBias and gcBias flags to correct for sequence-specific biases. We also used the recommended rangeFactorizationBins parameter value 4, which improves quantification accuracy on difficult-to-quantify transcripts. We then filtered out low-expression genes by excluding genes with a median expression of 0 within a dataset (SchildkrautB: 10,620 genes removed, SchildkrautW: 10,410 genes removed). We library-size normalized samples using upper quantile normalization. This normalization matches the 85^th^ percentile across samples to correct for library size differences across samples.

### Overview of comparator study populations

Consistent with the data processing pipeline used by Way *et al.* (7), we included data from the TCGA (platform: AffymetrixHT_HG-U133A), Yoshihara (platform: AgilentG4112F), and Tothill (platform: AffymetrixHG-U133Plus2) datasets included in the R package curatedOvarianData (21), and additional data from the dataset “Mayo” (GEO Accession: GSE74357, platform: AgilentG4112F). All gene expression data generated from these studies was derived from fresh frozen tumor tissue. Using the R package doppelgangR (22), we identified four more duplicate pairs across studies than found previously (7) and one pair of duplicate samples in SchildkrautW.

### Clustering Methods

We performed clustering analyses by modifying the pipeline from Way *et al.* (7) for inclusion of RNA-Seq data. Briefly, for K-means clustering, we used the R package cluster (version 2.0.6) (23), and for non-negative matrix factorization (NMF), we used the R package NMF (version 0.20.6) (24). We identified the top 1,500 genes with the largest Median Absolute Deviance (MAD) from each of the six datasets (SchildkrautB, SchildkrautW, TCGA, Mayo, Yoshihara, and Tothill), then used their union set of 4,355 genes for clustering (**Supplemental Table 1**). Consensus clusters for both K-means and NMF were identified using multiple random starts for each value of K=2-4 (K-means starts: 100; NMF starts: 10). Each dataset was clustered independently using the same set of genes.

We compared the similarity of clusters across datasets and clustering methods using the same approach as Way *et al.* (7). We used significance analysis of microarray (SAM) (25,26) to get cluster-specific vectors of moderated t-scores; this provided us with gene-specific expression patterns of each cluster within each dataset. We restricted to genes that were assayed and expressed across all datasets, which we denote as “common genes’’ (**Supplemental Table 2**).

Using these 8,360 “common genes’’, we compared clusters across: (1) datasets, and (2) clustering methodologies by calculating the Pearson correlation between two vectors of moderated t-scores. To use this approach on the RNA-Seq datasets, we log10(x+1) transformed the normalized counts to more closely match the data distribution of the microarray datasets. We performed a simulation study to examine SAM’s performance on log10 transformed counts relative to DESeq2’s performance on raw counts and found comparable performance (**Supplemental Note 1**).

Additionally, we compared the similarity of clusters in a global and aligned principal components analysis (PCA) projection. We filtered the datasets to only include the MAD genes used in our clustering pipeline, then within-sample scaled the expression values to ensure comparative ranges of expression across all datasets. We log10(x+1) transformed the RNA-Seq expression values before scaling to better match the RNA-Seq and microarray data distributions.

After normalization, we concatenated the datasets to calculate a global PCA projection. We then subtracted each dataset’s total centroid from each dataset’s cluster centroid to align each dataset such that it was centered at the origin of the PCA projection. This allowed us to compare the relative differences between each cluster across all datasets.

### Survival analyses by cluster

We generated Kaplan-Meier survival curves to visualize overall survival and subtype-specific survival within each dataset. We used Cox proportional hazards (PH) models to calculate hazard ratios (HR) and 95% confidence intervals (CIs) quantifying the differences in survival across HGSC subtypes. For five of the six datasets (SchildkrautB, SchildkrautW, TCGA, Mayo, Tothill), Cox PH models were adjusted for three factors that independently influence survival: age (in 5-year age groups), stage (I, II, III and IV), and debulking status (optimal debulking, suboptimal or missing debulking status). Yoshihara *et al.* did not provide information on age at HGSC diagnosis, so models for this dataset were adjusted for stage and debulking status only.

### Cluster comparison between pipeline runs and consensusOV

To ensure that our run of the updated Way *et al.* (7) pipeline was producing similar results to those from previously published subtype predictors, we compared the assigned cluster labels for each case in the TCGA, Mayo, Yoshihara, and Tothill datasets. To quantify the similarity between pipeline results, we rephrased the cluster comparison problem as a prediction problem. The ground-truth labels were the previously published cluster IDs (7) or consensusOV (27) predictions for each case, and the predicted labels were our newly assigned cluster labels. consensusOV version 1.16.0 was run using default parameters and no significance filtering. We used balanced accuracy as the metric for comparing current and published labels due to variations in cluster sizes. Balanced accuracy is defined in the following equations.

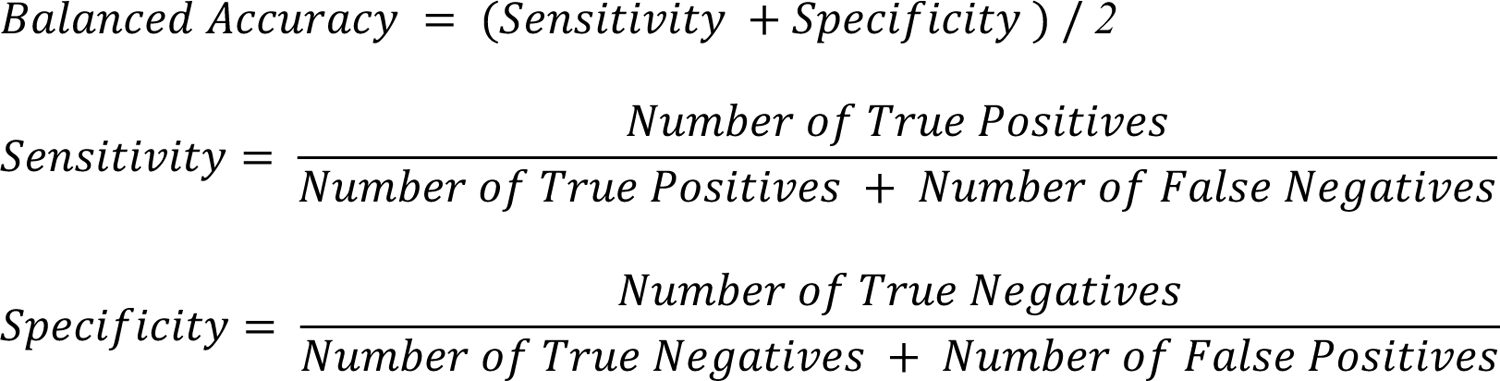

### Sensitivity analysis of K-means cluster assignments

K-means is sensitive to outliers and local minima; therefore, we quantified the stability of the cluster labels when using a subsample within each dataset. First, we removed any samples that were more than 1.5x the interquartile range from the first or third quartiles in the first five PCs. Using the remaining samples, we subsampled 80% of the samples within each dataset and re-ran K-means clustering. We then matched the new cluster labels to the original cluster labels using a greedy approach. We matched labels by finding the cluster IDs with the highest sample overlap. We did this iteratively, removing each cluster from consideration once its matched cluster ID was assigned.

### Data Availability Statement

We provide all software under the BSD 3-Clause License. We include scripts for the analyses presented in this manuscript, including RNA-Seq quantification, quality control, clustering, and figure generation. To support reproducibility, we provide code to recreate the environments and re-run both the RNA-Seq data analysis (https://github.com/greenelab/hgsc_rnaseq_cluster/) and our updated clustering pipeline (https://github.com/greenelab/hgsc_rnaseq_clustering_pipeline/). All derived expression data are publicly available and included in Supplemental Tables 6 and 7.

## Results

### Data quality

For both SchildkrautB and SchildkrautW, we further excluded samples due to possible technical artifacts (**Supplemental Tables 3 and 4**). In SchildkrautB and SchildkrautW, 14 and 4 cases were re-sequenced, respectively, due to insufficient read depth, and the transcriptomic profiles from the first sequencing attempt were excluded. After resequencing, all18 cases attained a read depth comparable to the other sequenced samples and progressed to the next step of the quality control pipeline (**Supplemental Figures 1B** and 2B). All other samples in both datasets were considered high quality, with over 83.3% of reads having a base quality of at least 30 (**Supplemental Figure 3**). After normalization, read count distributions were similar across sequencing batches, including samples previously identified by the sequencing core as low-quality (**Supplemental Figure 4C-H)**. However, a small subset of samples had lower-than-expected read counts. To account for this, we removed all samples where the median normalized read count was below 925 or where the bottom 25th quantile read count for a sample was below 30. This additional read count filter removed 10 SchildkrautB cases and 5 SchildkrautW cases (**Supplemental Figure 4A-B**). The final technical exclusion for SchildkrautB and SchildkrautW removed samples flagged by doppelgangR (22) as having overly similar expression patterns, suggesting they originated from the same tumor. Only SchildkrautW had a pair of samples identified as too similar (NCO0557 and NCO0625).

Following the SchildkrautB and SchildkrautW technical exclusions, we applied the Way pipeline exclusions to the TCGA, Mayo, Yoshihara, and Tothill datasets, and we applied doppelgangR exclusions to all datasets. The sizes of our final analytic data sets were as follows: SchildkrautB (n=262); SchildkrautW (n=309); TCGA (n = 499; <u>phs000178</u>) (5); Mayo (n = 377; <u>GSE74357</u>) (9); Yoshihara (n = 255; <u>GSE32062.GPL6480</u>) (28), and Tothill (n = 241; <u>GSE9891</u>) (6).

### Validation of updated clustering pipeline

We updated an HGSC subtyping pipeline (7) that previously identified clusters across four microarray studies from the United States, Japan, and Australia. While most of our pipeline was the same, we used balanced accuracy to quantify the similarity in output between pipelines. Balanced accuracy for our K-means clusters was consistently ≥0.91 across all datasets for K=2-4 (**Supplemental Figure 5**). The limited differences that arose were primarily due to changes in stochastic elements of the clustering methods and software packages, such as differences in computing environments, random seeds, and package versions. Another source of variation between the output of the pipelines was the addition of the Schildkraut datasets since the genes used for clustering must be shared across all datasets and have a high MAD in at least one dataset (**Methods)**. After adding the Schildkraut datasets, 893 genes that were previously denoted as common were reclassified as MAD genes, 609 MAD genes were re-classified as common genes, and 911 genes were removed from consideration. A Venn diagram of the removed genes is provided in **Supplemental Figure 6**.

To ensure that clustering outcomes were robust across methods, we compared the clusters from two different clustering methods: K-means and NMF. We derived a gene expression pattern for each cluster using the significance analysis of microarray (SAM) moderated t-score (**Methods**). Using this cluster-specific gene expression pattern, we calculated the Pearson correlation between moderated t-scores for clusters identified by each method. We found extremely high cluster concordance in all datasets between the clusters identified by NMF and K-means when we selected two and three clusters. We saw diminished concordance between methods when we used four clusters. Clusters in all datasets (except the Mayo dataset for K=4) had the highest concordance with the matched cluster along the diagonal (**Supplemental Figure 7**). Since K-means clustering is sensitive to outliers, we additionally ran K-means clustering in subsets of 80% of the data and found that, in all datasets, the majority of samples maintained their same cluster label in more than 50% of the subsampled reclusterings (**Supplemental Figure 8)**.

Our final clustering validation was to compare our output to output from another external HGSC subtype classifier, consensusOV (27). consensusOV combines four methods (5,6,8,9,29) into a consensus classifier. Since consensusOV can only assign samples to the four Verhaak-defined clusters, we only compare our K-means results for K=4. We found that our subtype calls were very similar to the consensusOV calls across all microarray datasets, with a minimum per-class balanced accuracy of 0.72 (**Supplemental Table 5**). We saw more discordance between our calls and consensusOV when we compared RNA-Seq samples, with per-class balanced accuracy ranging from 0.589-0.818 (**Supplemental Table 5**).

### Gene expression patterns in the self-identified Black study population compared with other study populations

Expression patterns for each cluster were summarized in two ways, first by plotting the principal components of normalized and aligned centroids in each dataset, and second, using SAM-moderated t-scores (7). In both approaches, we sought to determine if the relative cluster distances for each individual dataset were consistent across datasets. In our principal components (PC)-based approach, we normalized and projected all datasets into a shared PC space, then centered them at the origin of the space. For K=3, PC2 and PC3 could separate each cluster (**Figure 1a**). For K=4, PC2 and PC3 could not independently separate each cluster, but, together, could differentiate them (**Figure 1b**). Furthermore, in the first three PCs, the SchildkrautB dataset did not cluster away from any other datasets, and clustered closely with the SchildkrautW, TCGA, and Tothill datasets.

**Figure 1.**
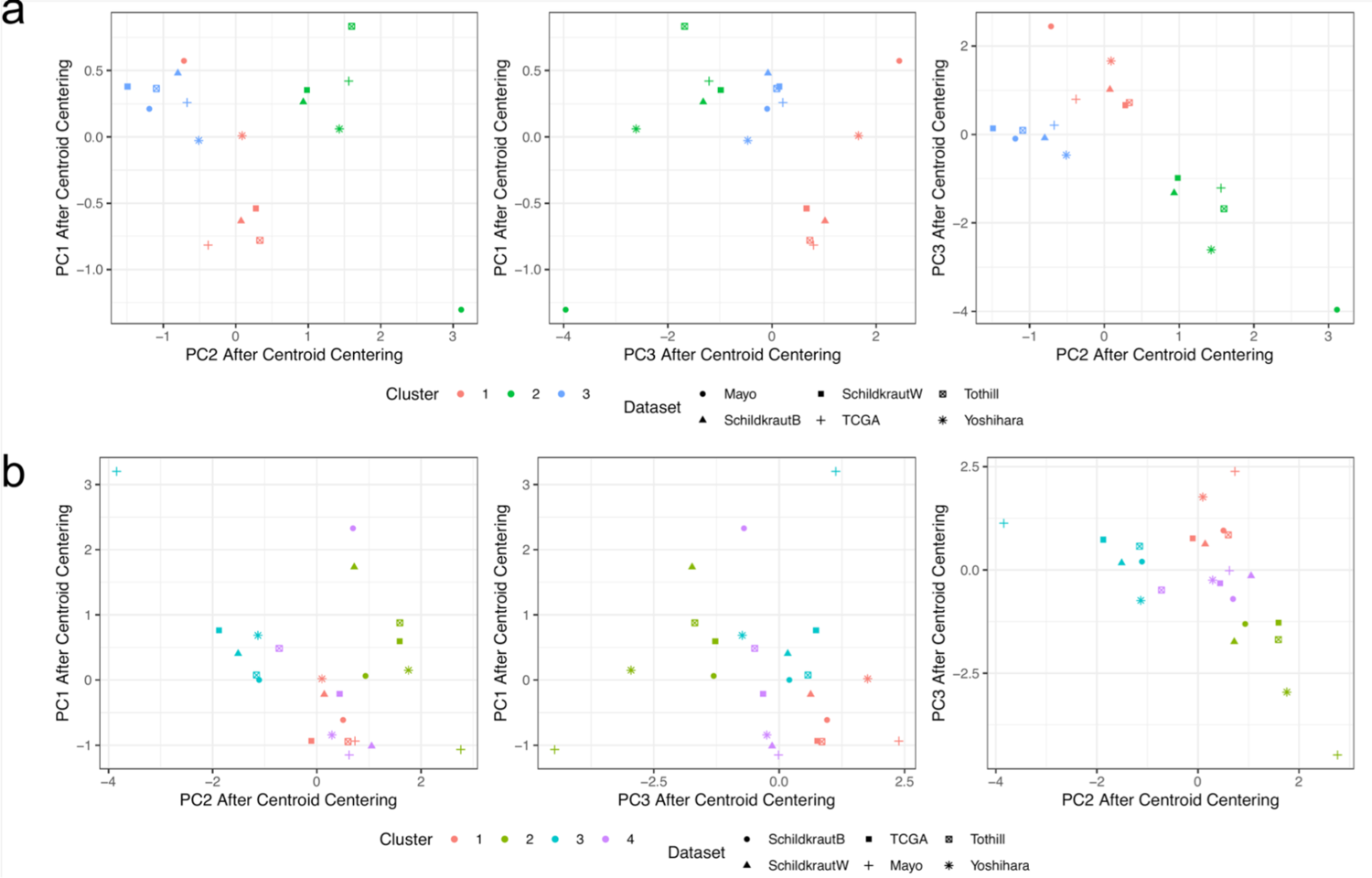
Principal Components Analysis (PCA) plots of cluster centroids for each dataset, after dataset alignment. **Panel a** compares K-means cluster centroids (K=3), and **Panel b** compares K-means cluster centroids (K=4) for each dataset considered in this study. We find that the principal components separate each cluster centroid in a consistent way across almost all datasets. For K=3, PC2 and PC3 are both able to separate each cluster independently, but for K=4 the combination of PC2 and PC3 are needed to separate each cluster. Furthermore, for K=3, we see that the Yoshihara and Mayo datasets have centroids that are much higher in PC1 than the other datasets. This trend continues for the Mayo dataset when K=4, in all PCs.

In our second, complementary approach, we compared differential gene expression between clusters by calculating the Pearson correlation for each cluster’s SAM-moderated t-score across pairs of datasets. **Figure 2** compares the cluster-specific gene expression pattern between the SchildkrautB cases and cases from the other five datasets. For K=3, SchildkrautB cluster-specific gene expression was highly correlated with cluster-specific gene expression across all datasets, and showed the strongest correlations with SchildkrautW cluster-specific gene expression. This provides strong evidence that the derived clusters from Black cases are the same as the derived clusters from White and Japanese cases. A high correlation between clusters was also found when performing pairwise comparisons among all six datasets (**Supplemental Figure 9**). Similar to Way *et al.* (7), we found that the correlation of cluster-specific gene expression patterns was diminished when using four, as opposed to three, clusters to describe the data. This was evident when comparing SchildkrautB to each of the five other populations (**Figure 2B** **vs.** **Figure 2A**) and when comparing gene expression patterns across all possible pairs of datasets (**Supplemental Figure 10 vs. Supplemental Figure 9**).

**Figure 2.**
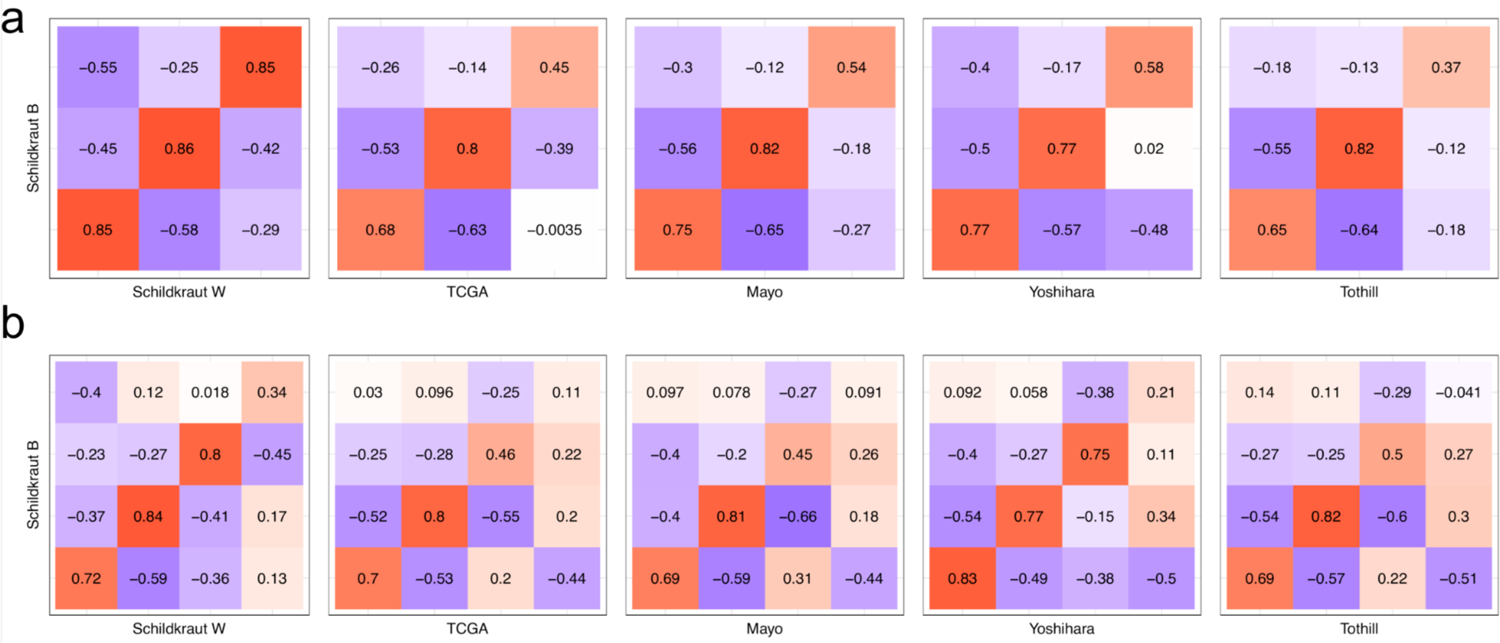
Significance analysis of microarray (SAM) moderated t-score Pearson correlation heatmaps of clusters across datasets. **Panel a** compares K-means clusters (K=3) between SchildkrautB and every other dataset considered in this study. Across each dataset we find a strong positive correlation with the clusters in SchildkrautB, with matched cluster correlations ranging from 0.37-0.86, and mismatched cluster correlations ranging from −0.65-0.03. **Panel b** performs the same comparison, but for K=4. In this comparison, we see much more inconsistency between matched clusters, with some mismatched clusters having a higher correlation than some matched clusters.

### Subtype distributions and characteristics by study population

We mapped the K-means clusters with K=4 to the four TCGA-derived HGSC subtypes to compare the frequency of subtypes across datasets and evaluate how cancer characteristics vary by subtype. The immunoreactive subtype was more common (39% vs 23-28%), and the differentiated subtype less common (7% vs 22-31%) in the self-identified Black study population when compared to the White and Japanese study populations (**Figure 3**). In analyses restricted to SchildkrautB and SchildkrautW, FIGO stage differed by HGSC subtype among both Black (*p*=0.040) and non-Hispanic White individuals (*p*=0.009); those with differentiated HGSC were more likely to have a lower FIGO stage at diagnosis (**Table 1**). Immune infiltration also differed by HGSC subtype, particularly among Black individuals (*p=*0.001 for CD3+ T cells; *p*=0.045 for CD3+/CD8+ suppressor T cells; **Table 1**). In both Black and White individuals, immune infiltration was higher for tumors categorized as mesenchymal or immunoreactive and lower for tumors categorized as proliferative.

**Figure 3.**
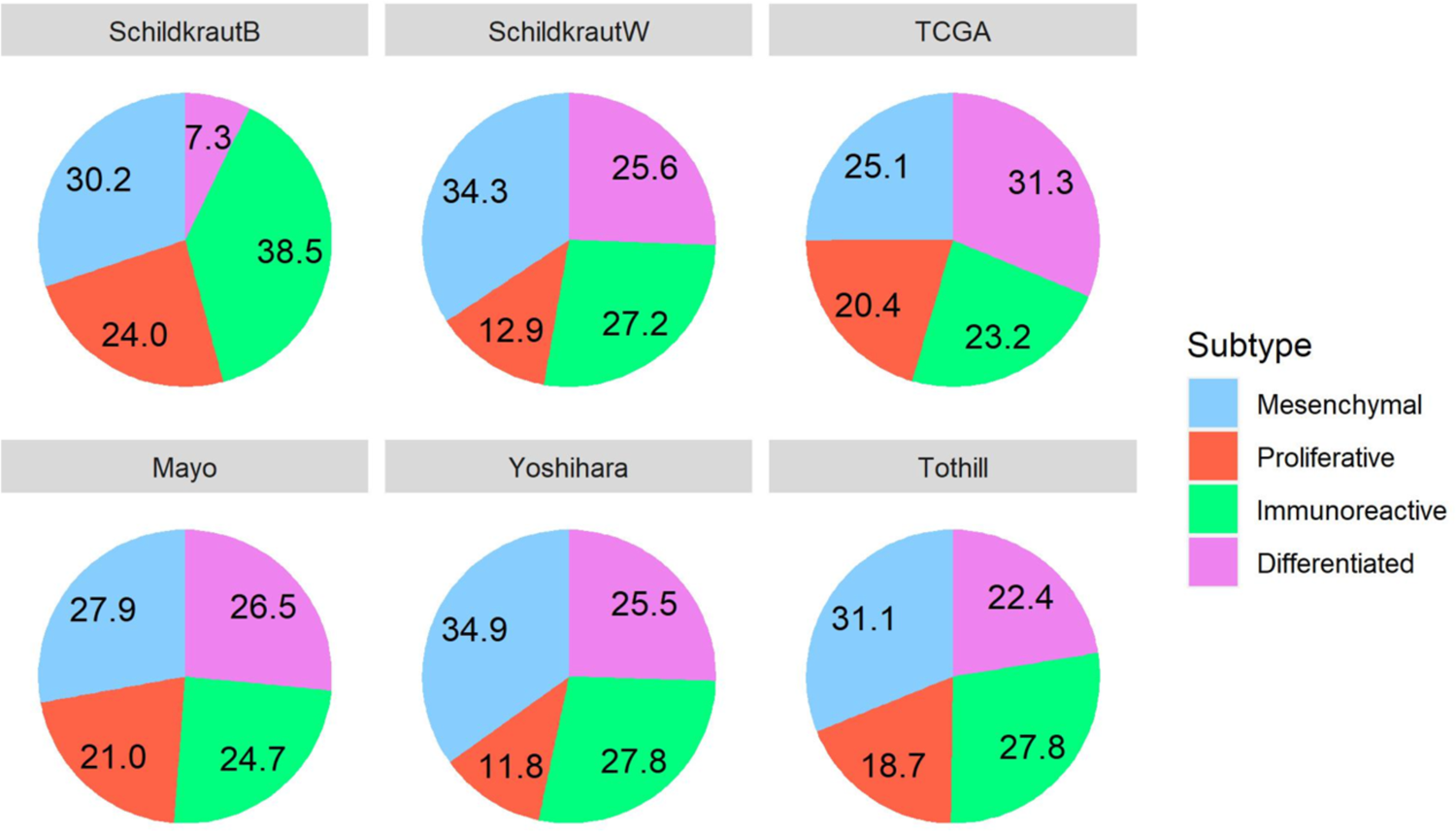
Distribution of subtypes across datasets.

**Table 1.**
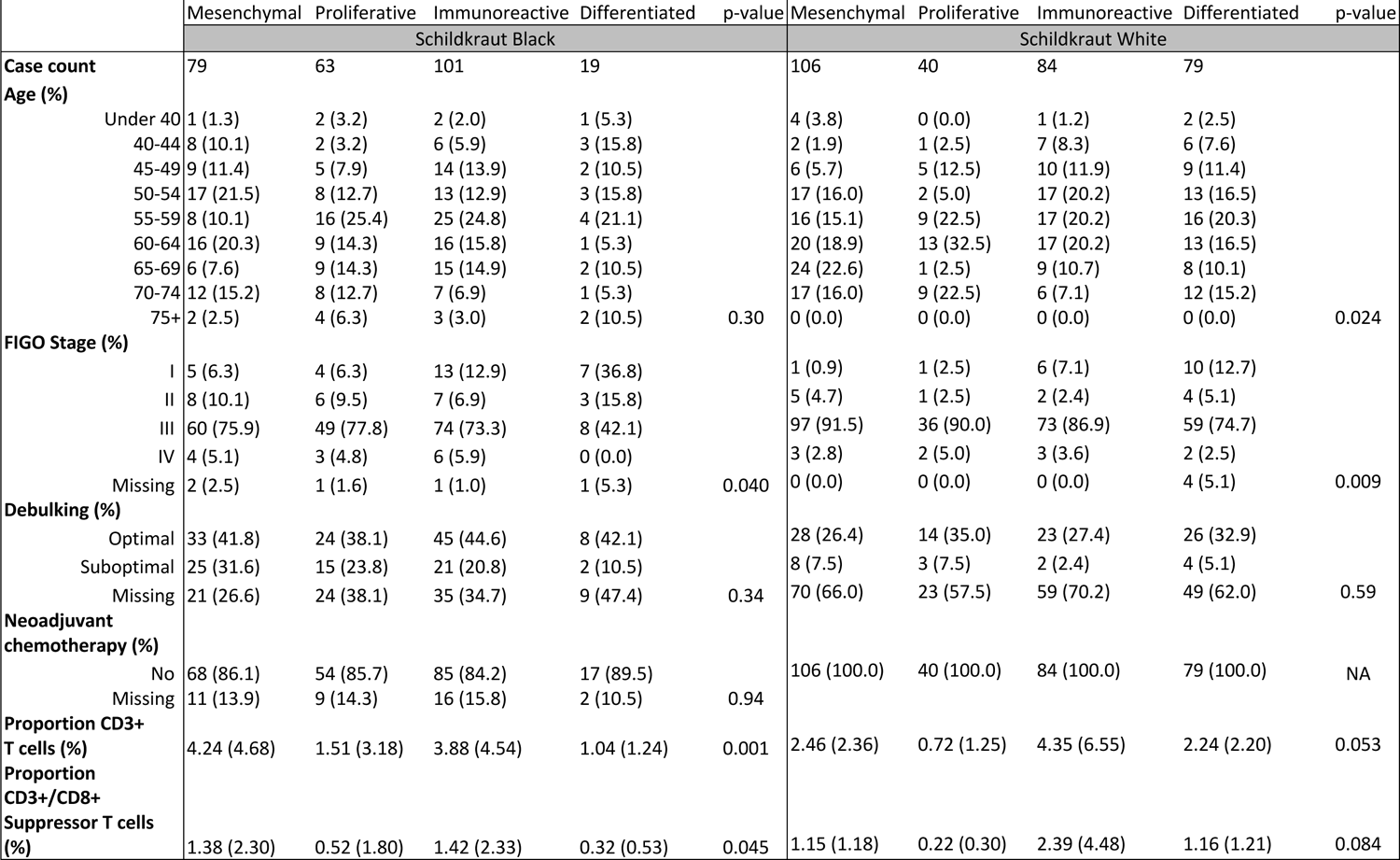
Characteristics of the Schildkraut study populations, by subtype.

### Survival patterns for clusters

Overall HGSC survival varied by study (**Supplemental Figure 11,** p-value for test of heterogeneity in a stage-adjusted model <0.001), though patterns of subtype-specific survival were generally similar across studies (**Figure 4**). Multivariable-adjusted hazard ratios and 95% CIs indicated that, when compared to individuals with mesenchymal tumors, those with immunoreactive tumors had better survival in most (SchildkrautB HR=0.79 [0.55, 1.13]; SchildkrautW HR=0.86 [0.62, 1.19]; Mayo HR=0.54 [0.39, 0.75]; Yoshihara HR=0.65 [0.39, 1.09]; Tothill HR=0.54 [0.31, 0.95]), but not all (TCGA HR=1.01 [0.69, 1.49]) study populations (**Table 2**). Meanwhile, the risk of death among those with proliferative HGSC was not statistically significantly different from the risk of death among those with mesenchymal tumors in any study population (**Table 2**), and it was close to the null value of 1.00 in both SchildkrautB (HR=0.98 [0.66, 1.46]) and SchildkrautW (HR=1.05 [0.70, 1.57]). The risk of death among those with differentiated HGSC was not statistically significantly different from the risk of death among those with mesenchymal tumors in five of the six study populations, and was lower in the Mayo study population (HR=0.61 [0.44, 0.84]; **Table 2**).

**Figure 4.**
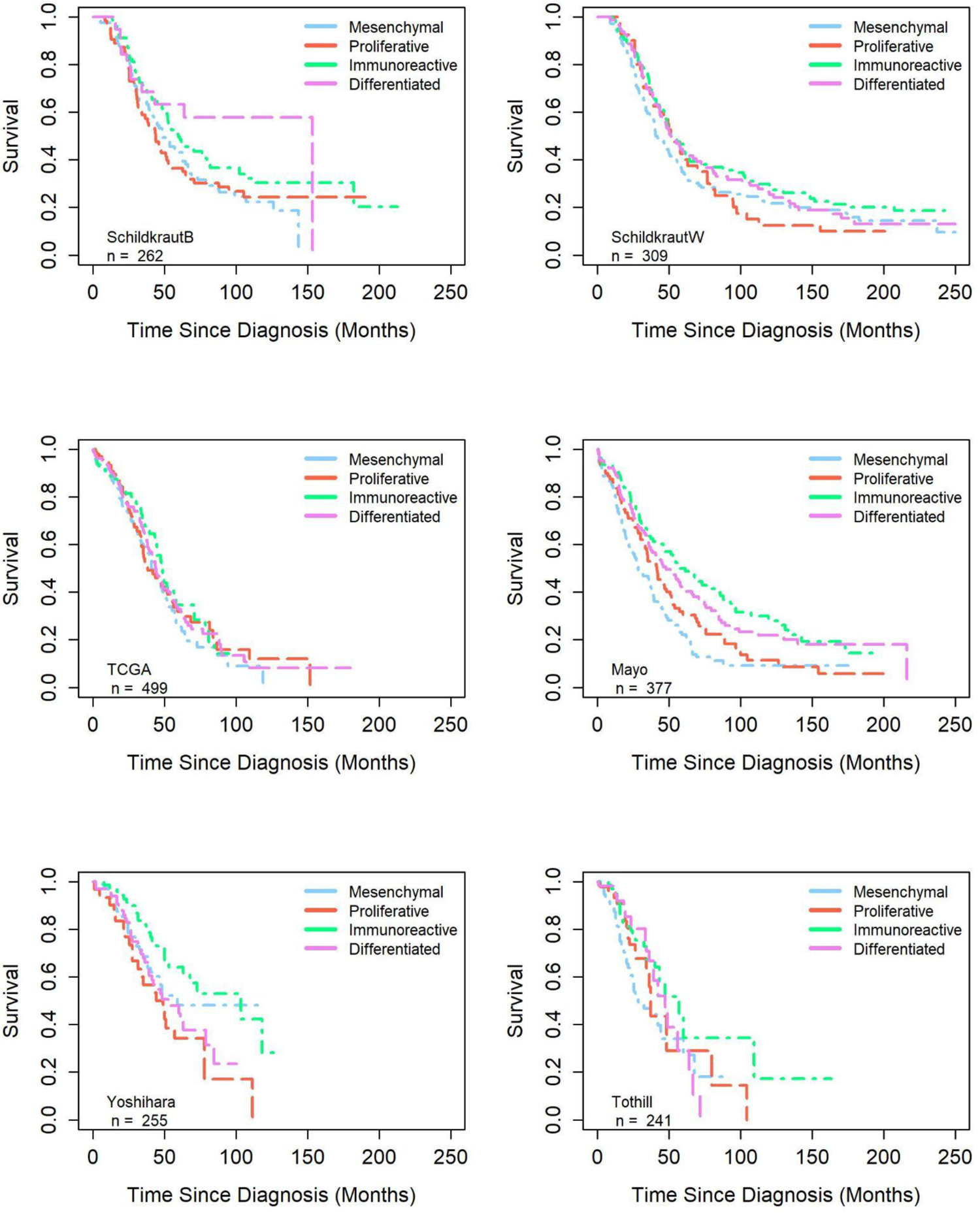
Kaplan Meier survival curves comparing subtype-specific survival by dataset.

**Table 2.**
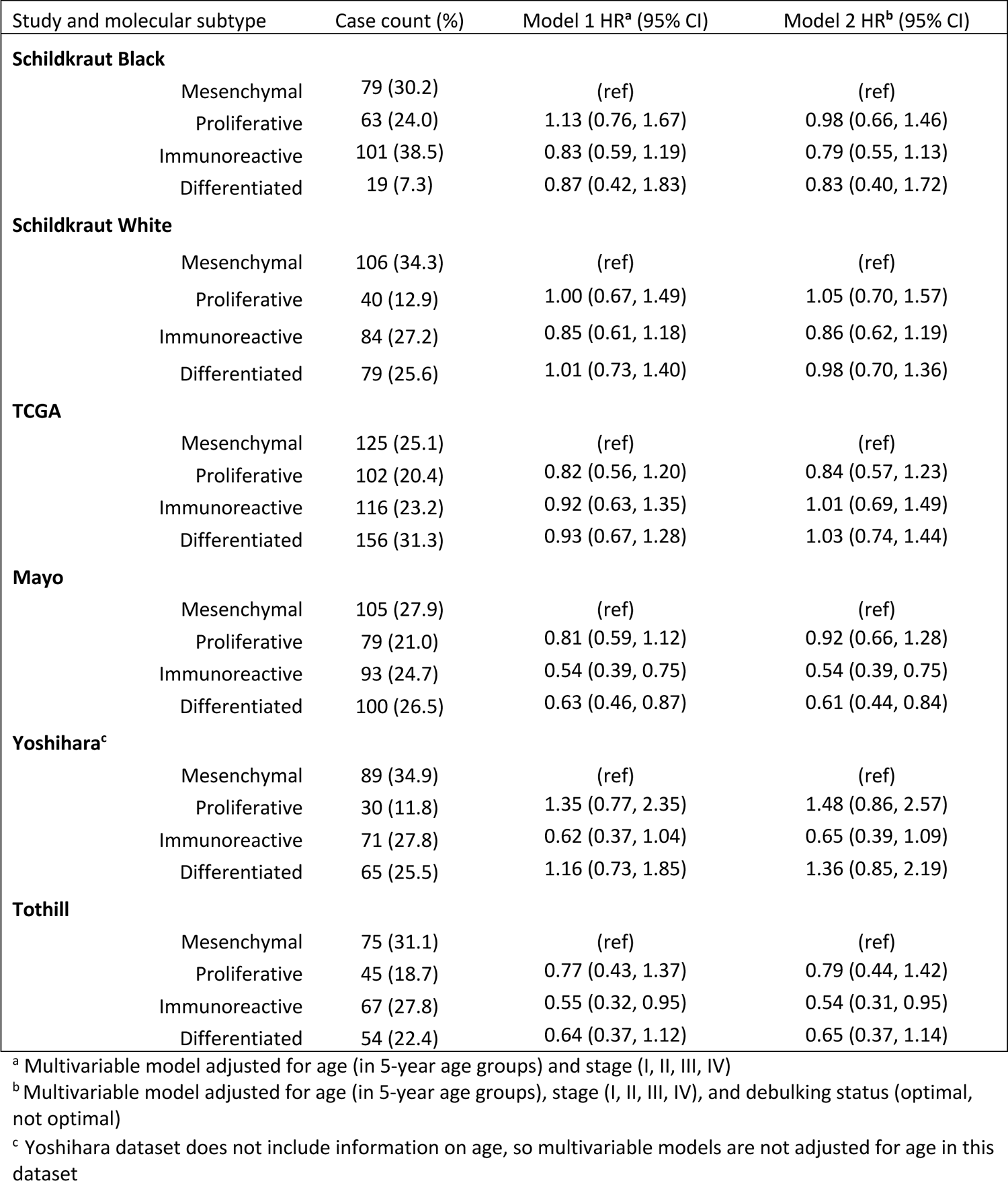
Subtype distribution and risk of death by study population.

| Study and molecular subtype | Case count (%) | Model 1 HR<sup>a</sup> (95% CI) | Model 2 HR<sup>b</sup> (95% CI) |
| --- | --- | --- | --- |
| **Schildkraut Black** |  |  |  |
| Mesenchymal | 79 (30.2) | (ref) | (ref) |
| Proliferative | 63 (24.0) | 1.13 (0.76, 1.67) | 0.98 (0.66, 1.46) |
| Immunoreactive | 101 (38.5) | 0.83 (0.59, 1.19) | 0.79 (0.55, 1.13) |
| Differentiated | 19 (7.3) | 0.87 (0.42, 1.83) | 0.83 (0.40, 1.72) |
| **Schildkraut White** |  |  |  |
| Mesenchymal | 106 (34.3) | (ref) | (ref) |
| Proliferative | 40 (12.9) | 1.00 (0.67, 1.49) | 1.05 (0.70, 1.57) |
| Immunoreactive | 84 (27.2) | 0.85 (0.61, 1.18) | 0.86 (0.62, 1.19) |
| Differentiated | 79 (25.6) | 1.01 (0.73, 1.40) | 0.98 (0.70, 1.36) |
| **TCGA** |  |  |  |
| Mesenchymal | 125 (25.1) | (ref) | (ref) |
| Proliferative | 102 (20.4) | 0.82 (0.56, 1.20) | 0.84 (0.57, 1.23) |
| Immunoreactive | 116 (23.2) | 0.92 (0.63, 1.35) | 1.01 (0.69, 1.49) |
| Differentiated | 156 (31.3) | 0.93 (0.67, 1.28) | 1.03 (0.74, 1.44) |
| **Mayo** |  |  |  |
| Mesenchymal | 105 (27.9) | (ref) | (ref) |
| Proliferative | 79 (21.0) | 0.81 (0.59, 1.12) | 0.92 (0.66, 1.28) |
| Immunoreactive | 93 (24.7) | 0.54 (0.39, 0.75) | 0.54 (0.39, 0.75) |
| Differentiated | 100 (26.5) | 0.63 (0.46, 0.87) | 0.61 (0.44, 0.84) |
| **Yoshihara<sup>c</sup>** |  |  |  |
| Mesenchymal | 89 (34.9) | (ref) | (ref) |
| Proliferative | 30 (11.8) | 1.35 (0.77, 2.35) | 1.48 (0.86, 2.57) |
| Immunoreactive | 71 (27.8) | 0.62 (0.37, 1.04) | 0.65 (0.39, 1.09) |
| Differentiated | 65 (25.5) | 1.16 (0.73, 1.85) | 1.36 (0.85, 2.19) |
| **Tothill** |  |  |  |
| Mesenchymal | 75 (31.1) | (ref) | (ref) |
| Proliferative | 45 (18.7) | 0.77 (0.43, 1.37) | 0.79 (0.44, 1.42) |
| Immunoreactive | 67 (27.8) | 0.55 (0.32, 0.95) | 0.54 (0.31, 0.95) |
| Differentiated | 54 (22.4) | 0.64 (0.37, 1.12) | 0.65 (0.37, 1.14) |

<sup>a</sup> Multivariable model adjusted for age (in 5-year age groups) and stage (I, II, III, IV)
<sup>b</sup> Multivariable model adjusted for age (in 5-year age groups), stage (I, II, III, IV), and debulking status (optimal, not optimal)
<sup>c</sup> Yoshihara dataset does not include information on age, so multivariable models are not adjusted for age in this dataset

## Discussion

We performed a cross-platform and cross-population analysis of HGSC, including four existing datasets, plus newly generated RNA-Seq gene expression data from 262 Black and 309 White HGSC cases. When comparing RNA-Seq data from the 309 White cases to microarray gene expression data from the predominantly White populations that comprise the Tothill and TCGA study populations, we observed high cluster stability within each dataset when advancing from K=2 to K=4 (**Supplemental Figure 8**) and consistent cluster-specific gene expression profiles across datasets (**Supplemental Figures 9 and 10**). This indicates that similar HGSC gene expression clusters can be defined using combined data from array-based technologies in fresh frozen tissue and RNA-seq technologies in FFPE tissue. We also observed consistent cluster composition (**Figure 1**) and gene expression profiles when comparing Black HGSC cases to White and Japanese HGSC cases (**Figure 2**). This indicates that HGSC gene expression clusters are consistent across Black, White, and Japanese individuals, so it is unlikely that racial differences in HGSC gene expression patterns are a key driver behind poorer HGSC survival in Black populations.

Our interest in determining how an existing HGSC subtype clustering pipeline performs with RNA-Seq data was motivated by the increasing use of RNA-Seq technology to interrogate cancer gene expression (30). Previously published HGSC subtype clustering approaches were designed around array-based gene expression data (5–7,9,28,31), and while array-based and RNA-Seq technologies observe the same underlying biological processes, the data they produce follow different data distributions (32,33). This is most clear in the methodological differences between array-based and RNA-Seq differential gene expression methods (34–37). Here, we demonstrated that, despite the difference in data distributions generated by array-based and RNA-Seq technologies, the Way *et al.* (7) subtype pipeline built for array-based gene expression data can be applied to normalized and log-transformed RNA-Seq data. The high cluster correlations across all four array-based datasets using fresh frozen tissue and both RNA-Seq datasets using FFPE tissue indicate that the delineation of HGSC subtypes is agnostic to sequencing technology and fresh frozen versus FFPE tissue.

As done in previous studies, we assigned all cases to one of the four TCGA-derived HGSC subtypes; however, as was previously reported in Way *et al.* (7), we observed that cluster-specific gene expression was more concordant across datasets when clusters were assigned using K=2 or K=3 compared to K=4. This was most evident when comparing each dataset’s cluster-specific expression profiles to the cluster-specific expression profiles across all of the other datasets (**Figure 2, Supplemental Figures 9 and 10**). When considering K=3, the largest off-diagonal correlation we observed over all datasets was 0.07 (**Supplemental Figure 9**). In contrast, for K=4 we found much larger positive off-diagonal correlations for the clusters observed in the array-based TCGA, Mayo, Yoshihara, and Tothill datasets, with the largest per-dataset correlations ranging from 0.19 to 0.33 (**Supplemental Figure 10**). We also observed large positive off-diagonal correlations for the RNA-Seq-based SchildkrautB and SchildkrautW datasets, with the largest per-dataset correlations ranging from 0.17 to 0.34 (**Supplemental Figure 10**). Since cluster-specific gene expression was more concordant for K=2 and K=3 versus K=4 for all datasets, we encourage future studies to test whether a number other than four best represents HGSC subtypes. Our results are consistent with a model where either two HGSC gene expression axes (e.g., mesenchymal-like and immune) or a set of three HGSC subtypes (e.g., as derived using K=3) may more effectively describe the biological variation in HGSC gene expression than the four subtypes most commonly seen in the literature.

Beyond methodological advances, our work also provides strong evidence that HGSC subtypes can be reproduced among Black HGSC cases and that Black, White, and Japanese HGSC cases share similar subtype-specific gene expression profiles. At increasing values of K, we observed patterns in cluster composition among Black HGSC cases that were consistent with those observed among White and Japanese HGSC cases (**Figure 1, Supplemental Figure 8**).

Further, we observed strong correlations between cluster-specific gene expression in Black cases and cluster-specific gene expression in White and Japanese cases, especially when comparing clusters defined using K=2 and K=3 (**Figure 2**). Patterns of subtype-specific survival were also generally consistent across populations. When compared to mesenchymal HGSC, the risk of death was similar for proliferative and differentiated cases, and lower, but not statistically significantly lower, for immunoreactive cases both in SchildkrautB (HR=0.79 [0.55, 1.13]) and SchildkrautW (HR=0.86 [0.62, 1.19], **Table 2**, **Figure 4**).

The primary difference we observed when comparing HGSC subtypes in Black individuals (i.e., cases in SchildkrautB) to HGSC subtypes in all other study populations was that more Black HGSC cases had gene expression profiles consistent with the TCGA immunoreactive subtype (39% compared to 23%-28%) and fewer Black HGSC cases had gene expression profiles consistent with the TCGA differentiated subtype (7% compared to 22%-31%; **Figure 3**). Differences in the Tothill, C1-C6, gene expression subtype signatures for Black (n=29) versus White (n=156) ovarian cancers were observed previously, though in different proportions (17), so it is possible that there exists true variation in the proportion of HGSC subtypes for Black versus non-Black HGSC cases. However, it is also possible that variations in study design across the six study populations contributed to the different subtype distributions that we observed. For example, case-control studies like the AACES and NCOCS are unable to enroll cases with rapidly fatal HGSC (15). This could have skewed the observed subtype distribution for Black cases toward a greater proportion of less aggressive, immunoreactive tumors, and artificially inflated estimates of overall survival in the SchildkrautB study population (**Supplemental Figure 11**), as has been posited previously (15).

A key contribution of this study was that we were able to update a previously published subtype clustering pipeline to accept either array-based gene expression data or RNA-Seq data and validate our modifications. We also created the first large RNA-Seq dataset of HGSC in self-identified Black cases, consisting of 262 high-quality expression profiles. This dataset allowed us to compare the expression profiles of HGSC subtypes in Black cases against other study populations, and it provided an opportunity to evaluate differences in subtype frequency and survival in Black HGSC cases compared to non-Black HGSC cases. An important limitation of this study was that we lacked adequate data to explore whether the observed racial variation in the proportions of gene expression subtypes and survival outcomes was due to biological, sociodemographic, or access-to-care differences.

In summary, we have updated an existing HGSC gene expression subtype classifier to be compatible with both array-based gene expression data and RNA-Seq data. This advancement will facilitate reproducible HGSC subtyping for research purposes and is available for use in future studies. We have also demonstrated that the HGSC subtypes generated by our classifier generalize to racially diverse populations, and we have indicated that HGSC subtype-specific gene expression and subtype-specific survival are consistent across Black, White and Asian study populations. Given our findings, we expect that a clinical HGSC gene expression assay would benefit prognostication and treatment strategies similarly for women from multiple racial and ethnic backgrounds.

## Acknowledgements

We would like to acknowledge the AACES interviewers, Christine Bard, LaTonda Briggs, Whitney Franz (North Carolina) and Robin Gold (Detroit). We also acknowledge the individuals responsible for facilitating case ascertainment across the ten sites including: Jennifer Burczyk-Brown (Alabama); Rana Bayakly and Vicki Bennett (Georgia); the Louisiana Tumor Registry; Lisa Paddock and Manisha Narang (New Jersey); Diana Slone, Yingli Wolinsky, Steven Waggoner, Anne Heugel, Nancy Fusco, Kelly Ferguson, Peter Rose, Deb Strater, Taryn Ferber, Donna White, Lynn Borzi, Eric Jenison, Nairmeen Haller, Debbie Thomas, Vivian von Gruenigen, Michele McCarroll, Joyce Neading, John Geisler, Stephanie Smiddy, David Cohn, Michele Vaughan, Luis Vaccarello, Elayna Freese, James Pavelka, Pam Plummer, William Nahhas, Ellen Cato, John Moroney, Mark Wysong, Tonia Combs, Marci Bowling, Brandon Fletcher (Ohio); Martin Whiteside (Tennessee) and Georgina Armstrong and the Texas Registry, Cancer Epidemiology and Surveillance Branch, Department of State Health Services. We would also like to acknowledge the AACES investigators, Anthony J Alberg, Elisa V Bandera, Jill Barnholtz-Sloan, Melissa Bondy, Michele L Cote, Ellen Funkhouser, Edward Peters, Ann G Schwartz, Paul Terry, and Patricia G Moorman. This study would not have been possible without the efforts of the North Carolina Central Tumor Registry and all of the staff of the NCOCS. We also thank Christine Lankevich for her management of the data collection for the North Carolina Study and Rex C. Bentley for review of the pathology in the NCOCS. We would like to thank Rex C. Bentley and Ann M. Mills for the review of the pathology in the AACES.

