## Supplementary figures and images for "Molecular subtypes of high-grade serous ovarian cancer across racial groups and gene expression platforms"

### SupplementalFigure1.JPG

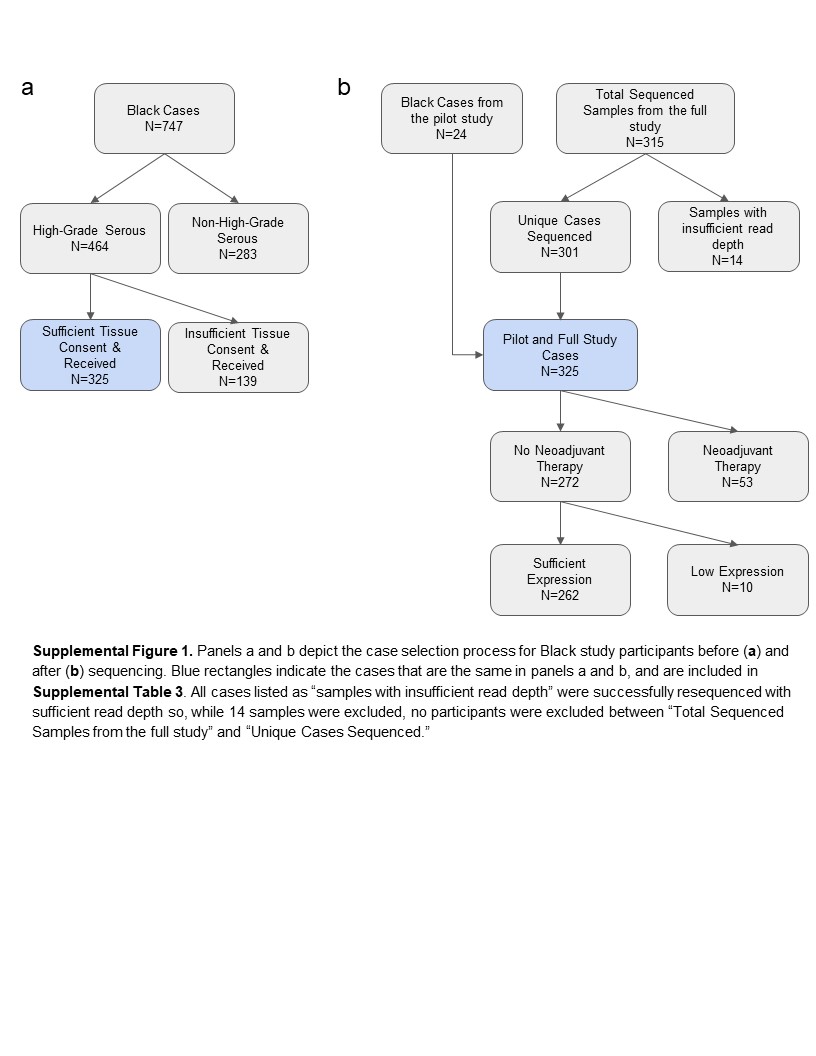

### SupplementalFigure2.JPG

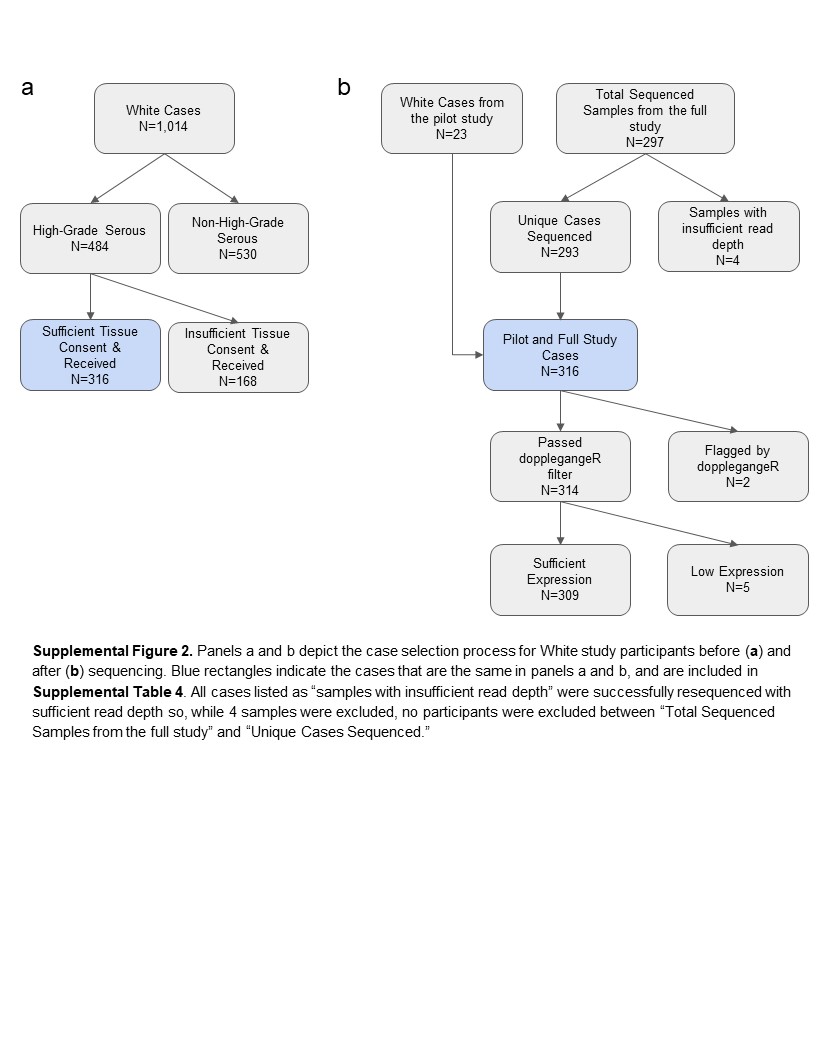

### SupplementalFigure3.JPG

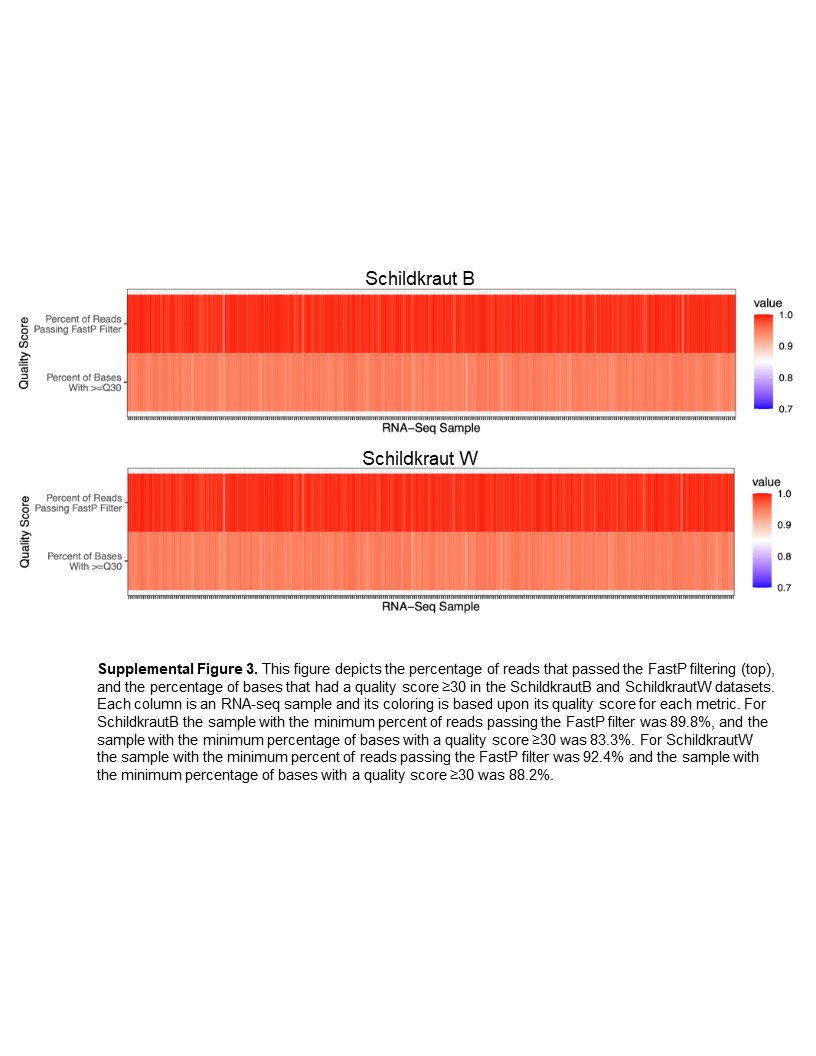

### SupplementalFigure4.JPG

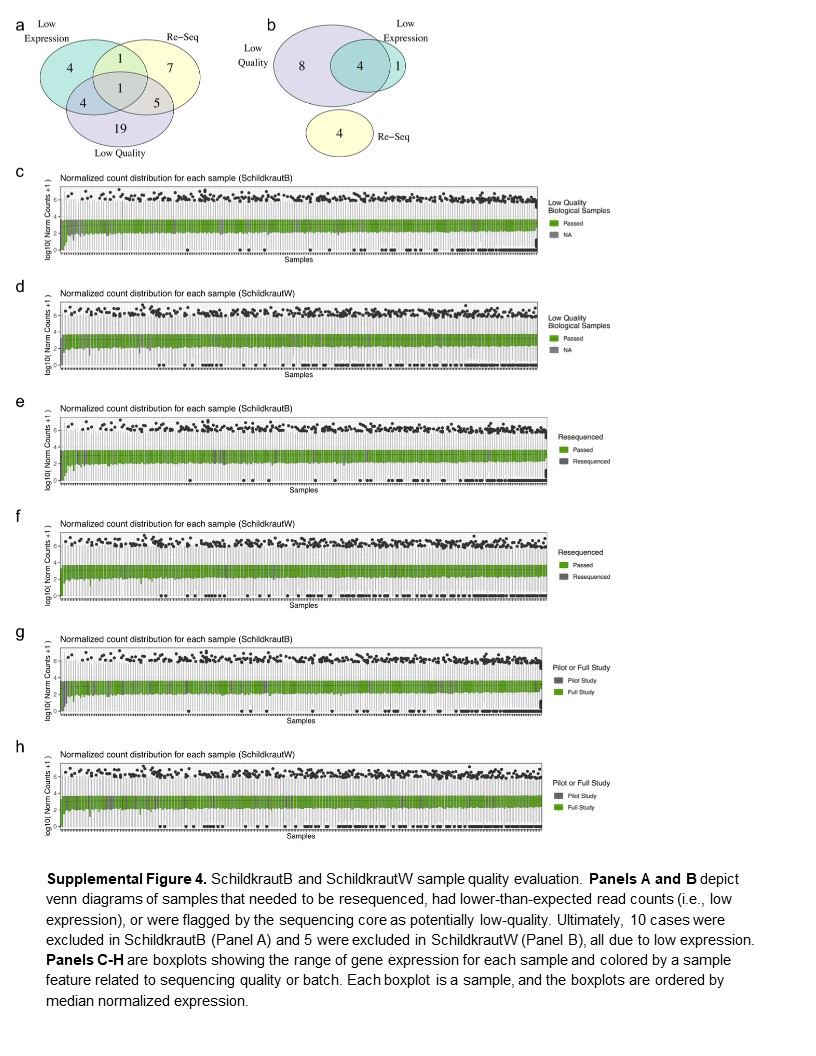

### SupplementalFigure5.JPG

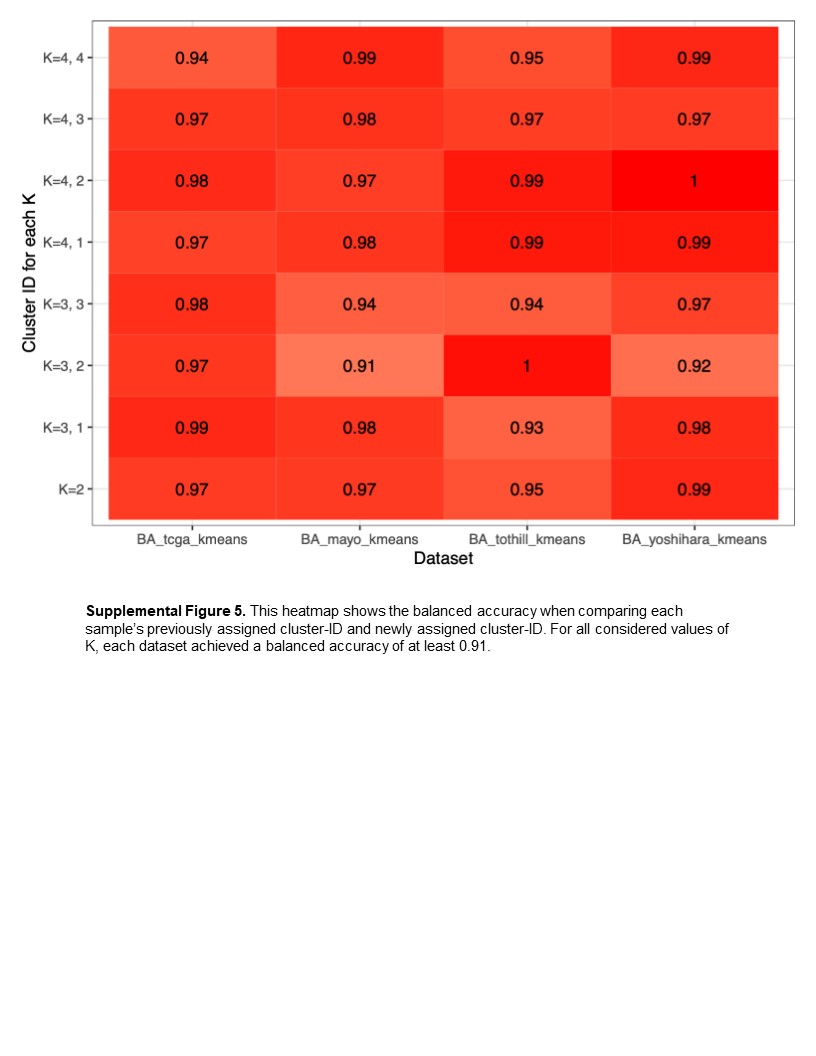

### SupplementalFigure6.JPG

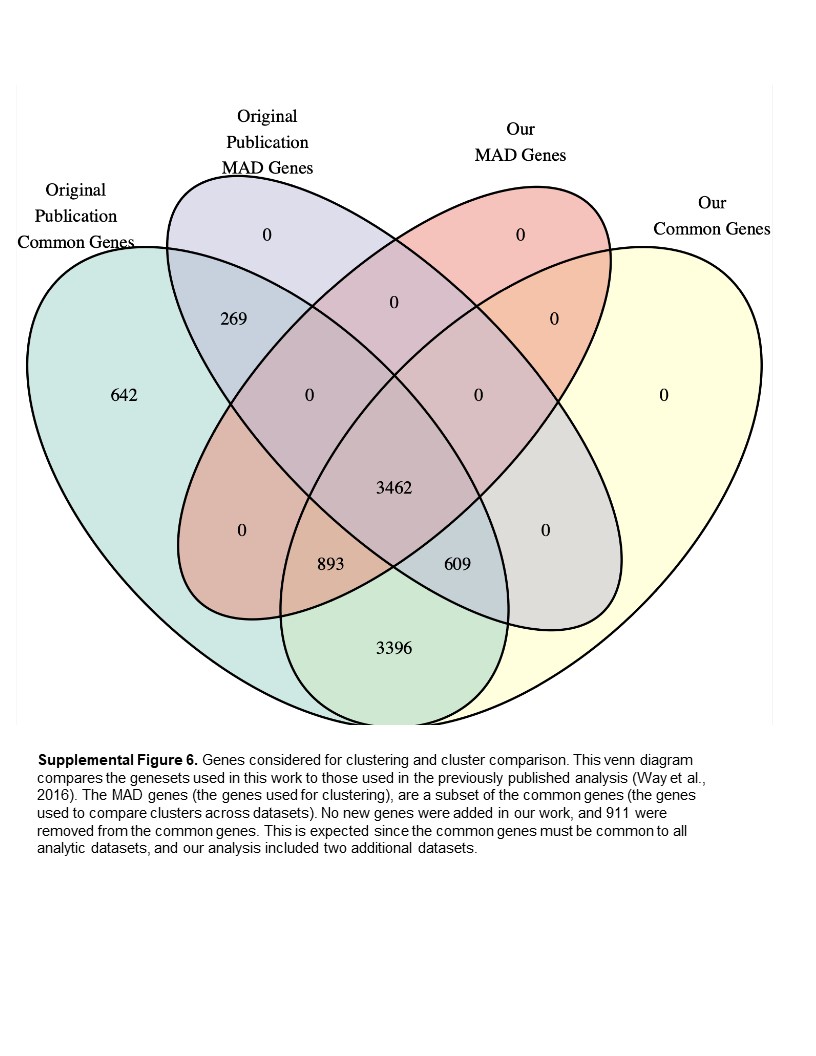

### SupplementalFigure7.JPG

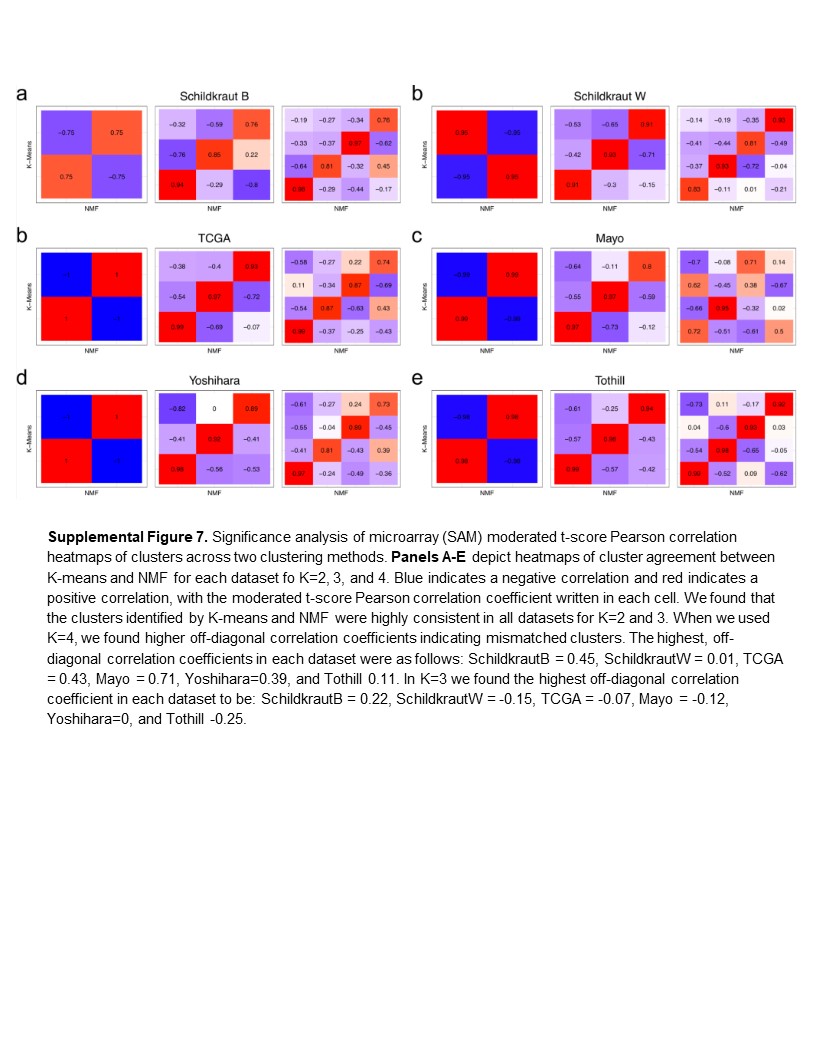

### SupplementalFigure8.JPG

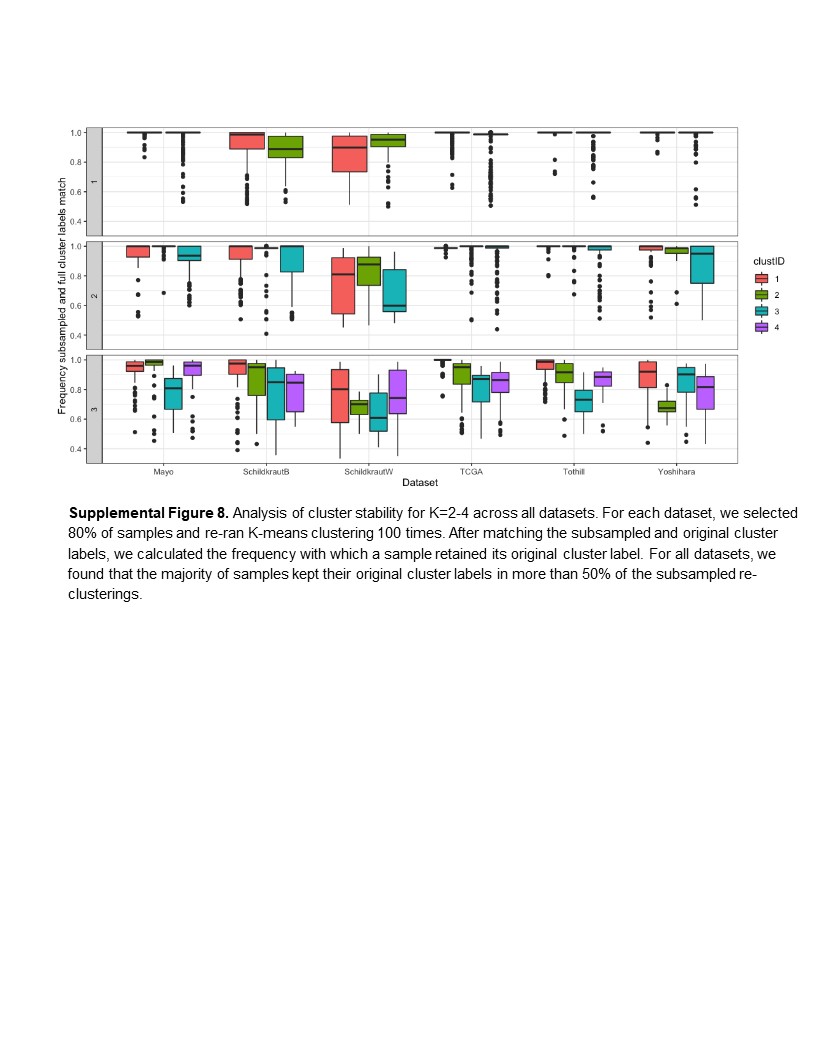

### SupplementalFigure9.JPG

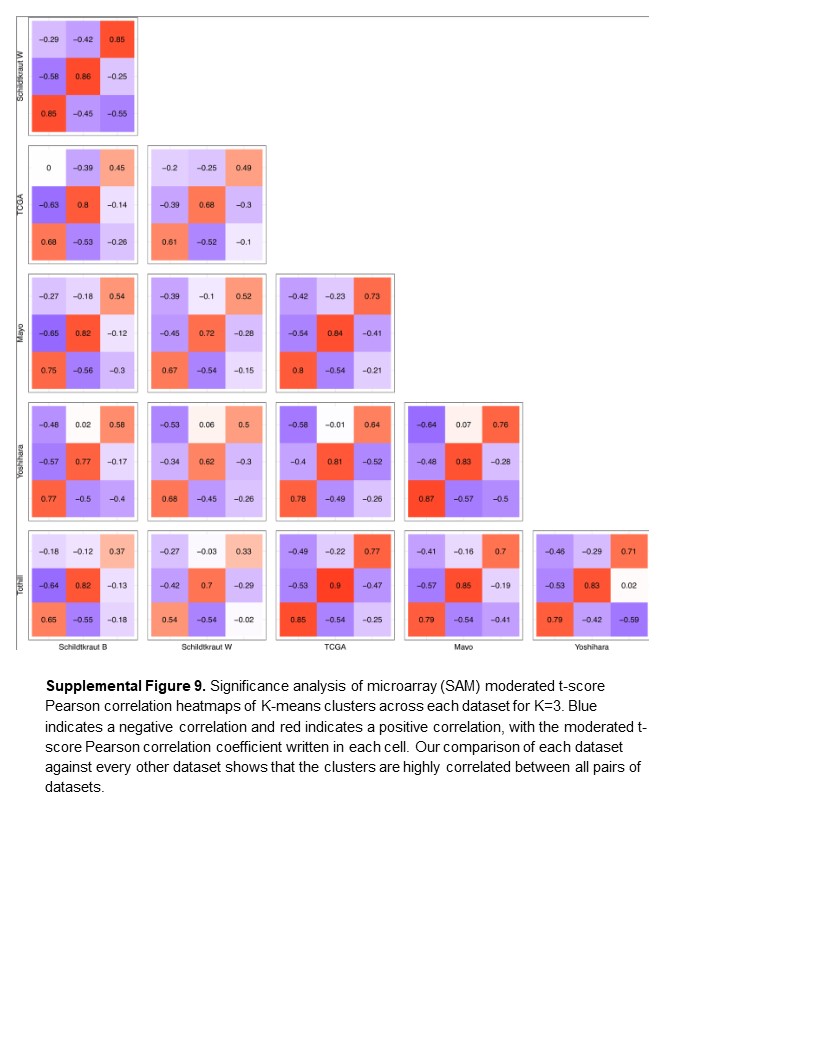

### SupplementalFigure10.JPG

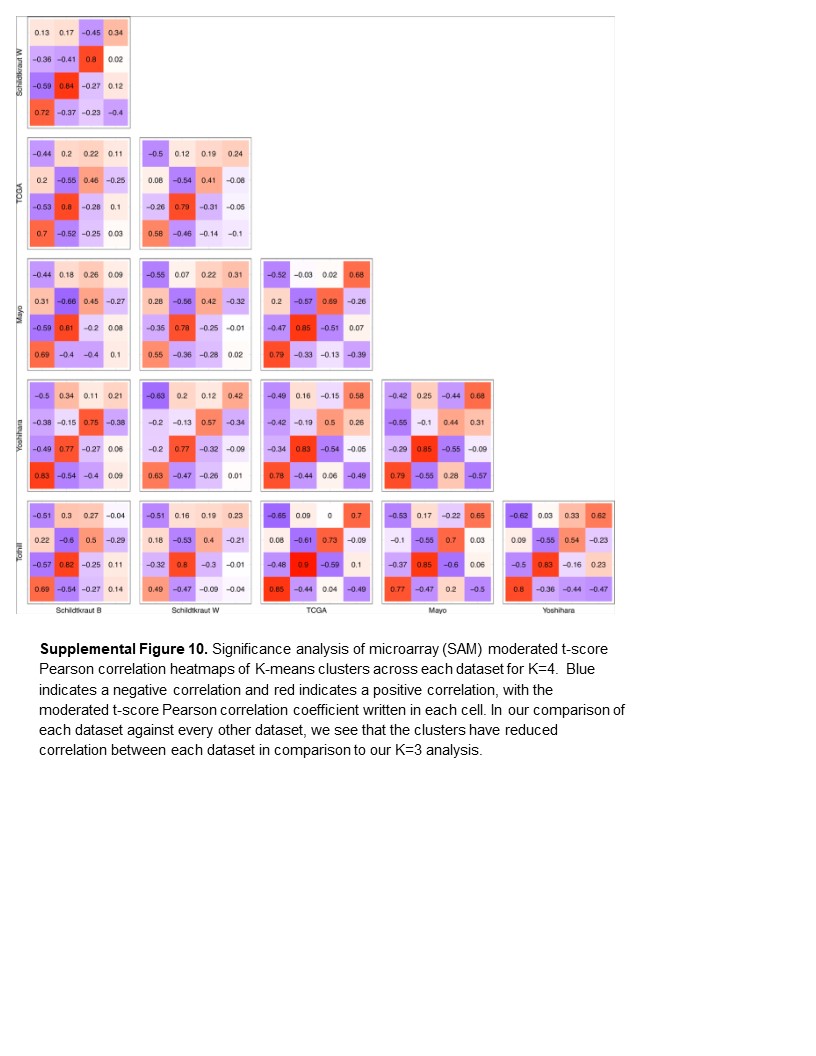

### SupplementalFigure11.JPG

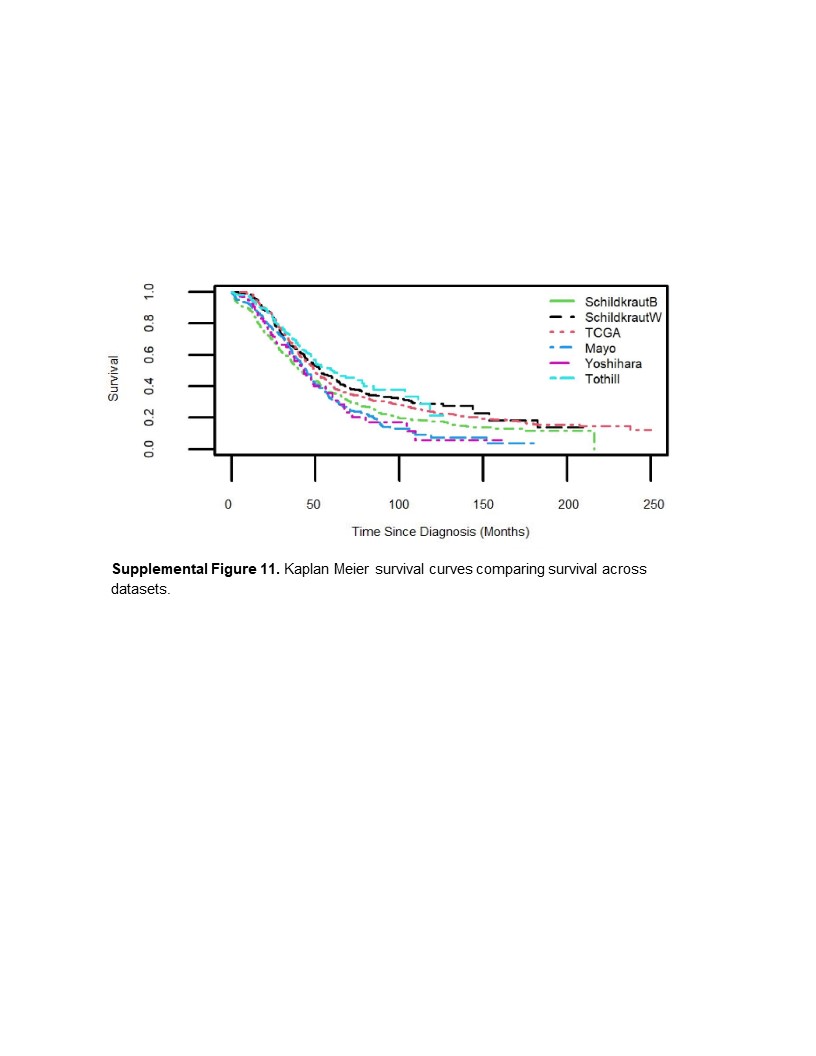
