## Supplemental Tables and Note for "Molecular subtypes of high-grade serous ovarian cancer across racial groups and gene expression platforms": supplemental_note_1.pdf

### test_SAM_performance

Natalie Davidson 22/06/2022

### run simulated DE on data from Way pipeline

#### get RNA-Seq data to be run in Way pipeline

The goal of this notebook is to see if the RNA-Seq data from the new AACES study can be run in the Way pipeline. Currently, the Way pipeline performs a differential expression test that identifies differentially expressed genes between clusters. The pipeline currently assumes microarray data and uses SAM to perform the test. So we will compare results using SAM and two other RNA-Seq specific methods. If SAM performs reasonably well in comparison to the other methods, we will continue using SAM.

Comparison will be done on:

1. SAM using log10(RNA-Seq counts)
2. EdgerR using normalized counts
3. DESeq using raw counts

#### simulate differential expression

We are using binomial thinning on our input data in order to simulate differential expression between two randomly partitioned groups.

```
## [1] 0.5000268
```

#### run SAM

Now run the differential expression test

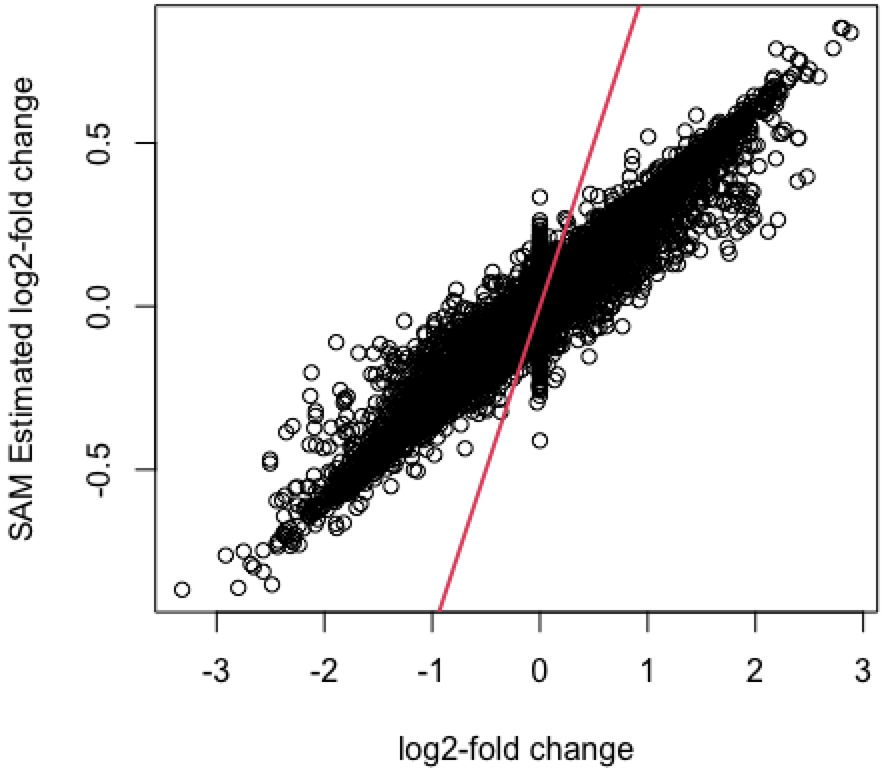
SAM Estimated log2-fold change
0.5
0.0
-0.5
-3
-2
-1
0
1
2
3
log2-fold change

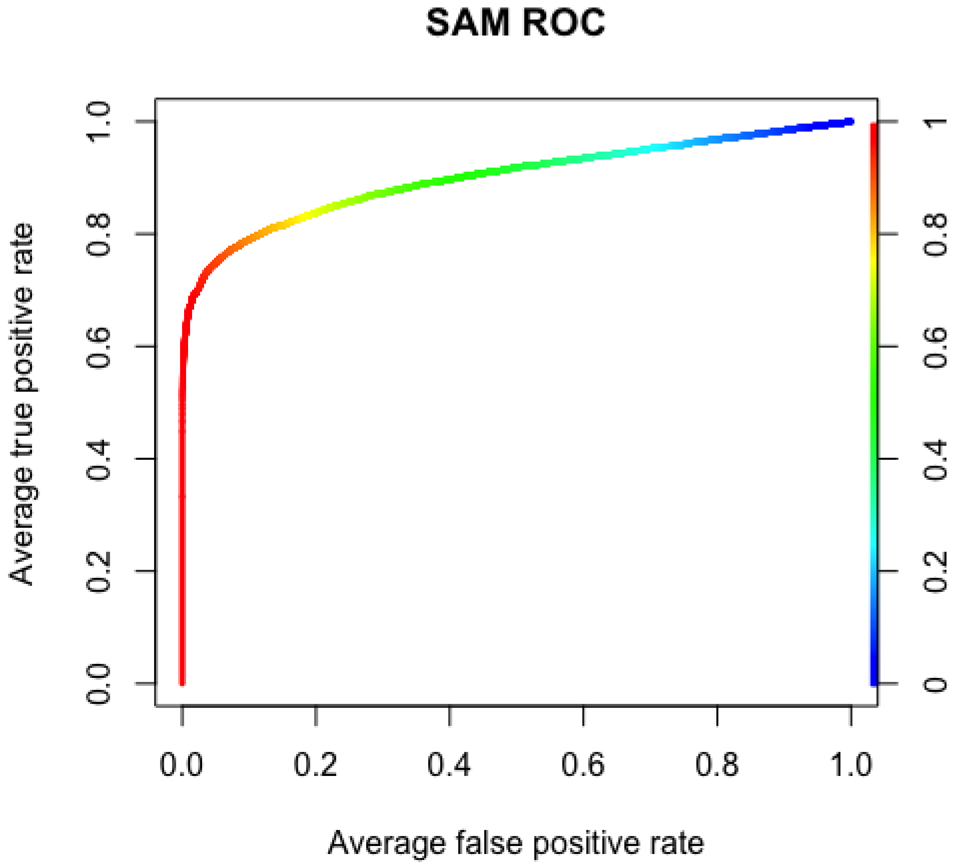
SAM ROC
Average true positive rate
0.0
0.2
0.4
0.6
0.8
1.0
0
0.2
0.4
0.6
0.8
1
Average false positive rate

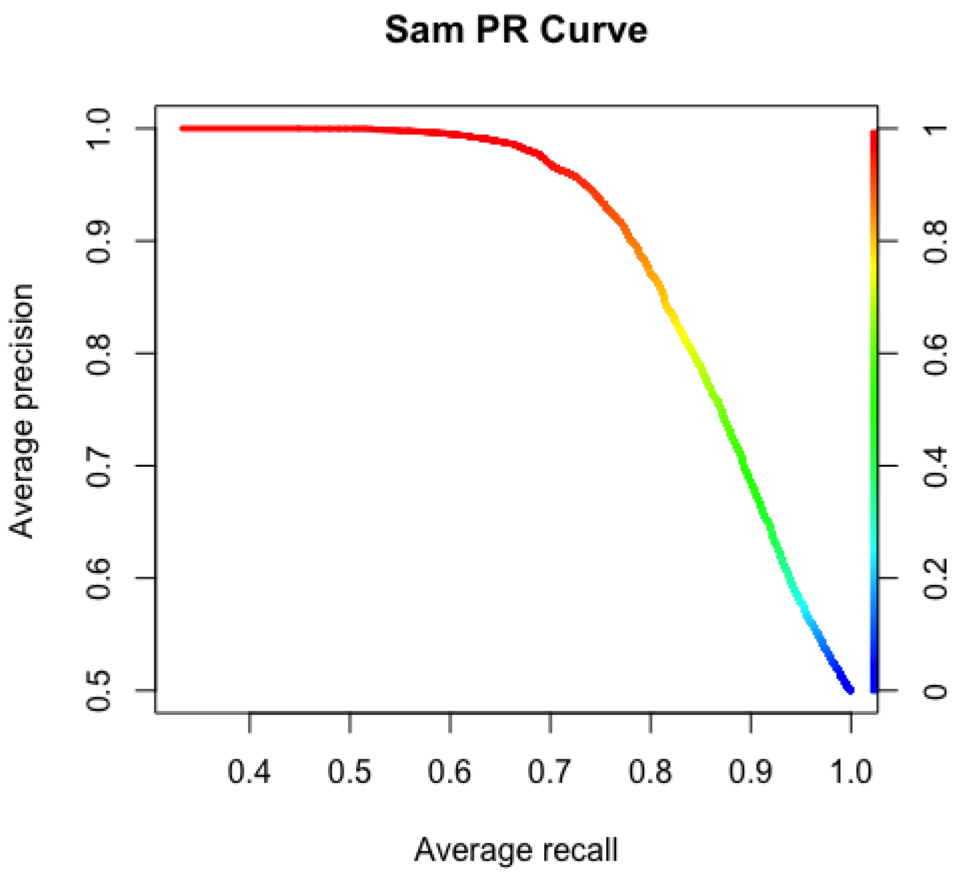


### check SAM

Since we have ground truth, report the performance

```
## Confusion Matrix and Statistics
##
##           Reference
## Prediction    0    1
##          0 9077  260
##          1 2694 6642
##
##                Accuracy : 0.8418
##                  95% CI : (0.8365, 0.847)
##     No Information Rate : 0.6304
##     P-Value [Acc > NIR] : < 2.2e-16
##
##                   Kappa : 0.6836
##
##  Mcnemar's Test P-Value : < 2.2e-16
##
##             Sensitivity : 0.9623
##             Specificity : 0.7711
##          Pos Pred Value : 0.7114
##          Neg Pred Value : 0.9722
##              Prevalence : 0.3696
##          Detection Rate : 0.3557
##    Detection Prevalence : 0.5000
##       Balanced Accuracy : 0.8667
##
##        'Positive' Class : 1
##
```

### make SAM QQ

Check test calibration

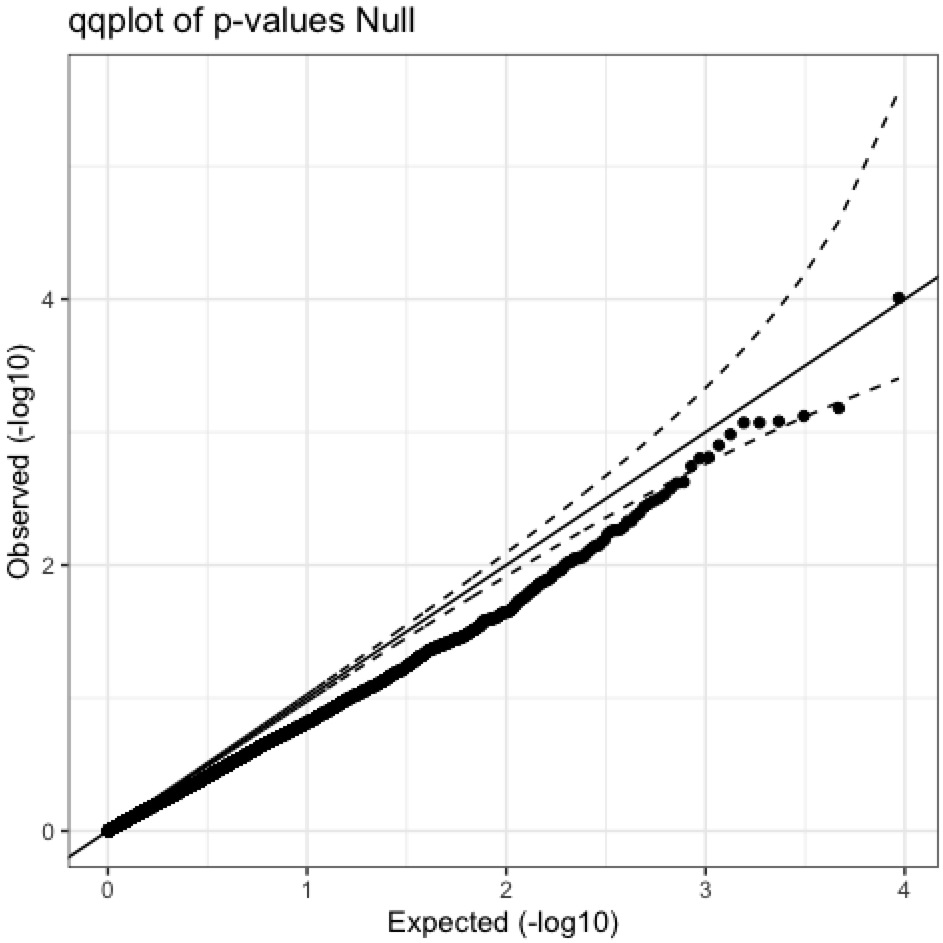


### run limma-voom

Now run the differential expression test using edgeR

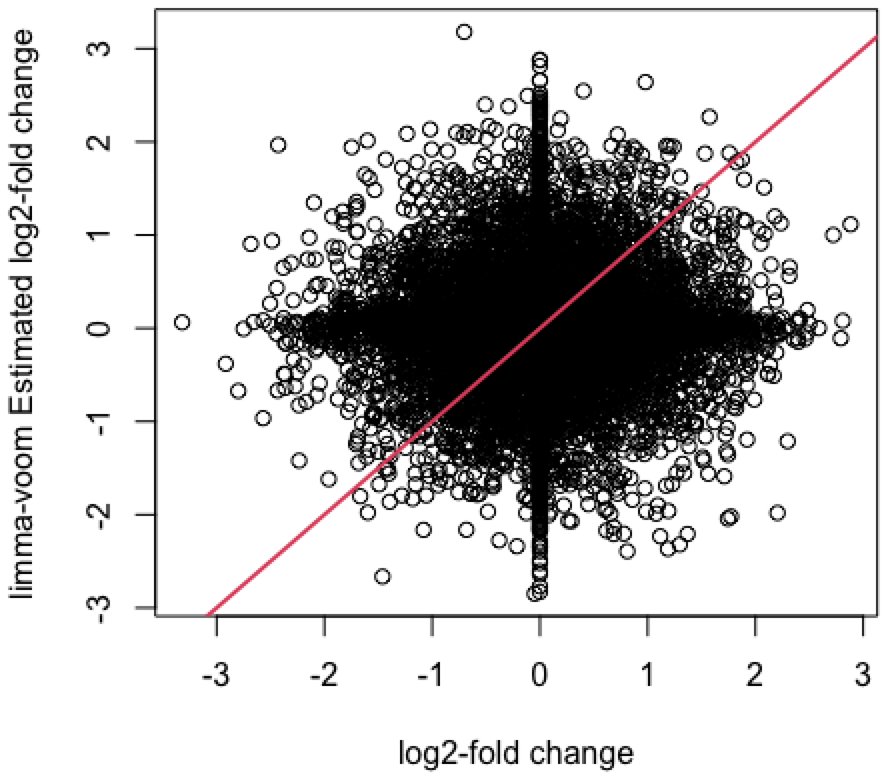
limma-voom Estimated log2-fold change
log2-fold change
-3
-2
-1
0
1
2
3

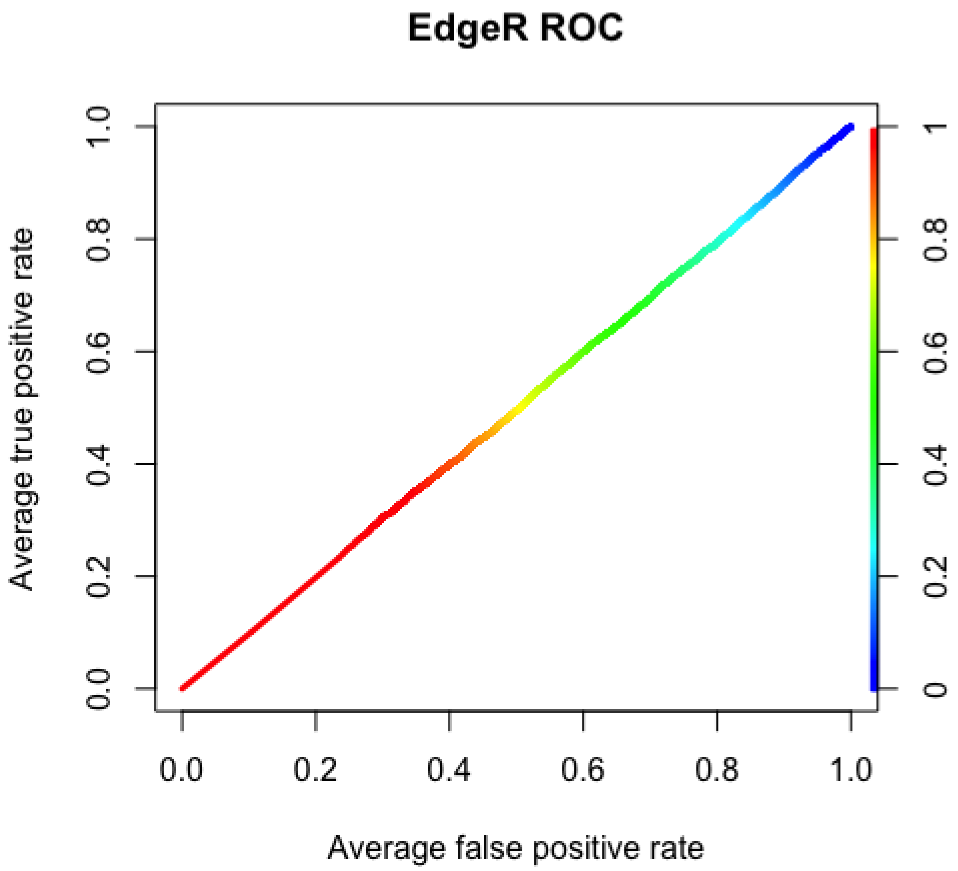
EdgeR ROC
Average true positive rate
Average false positive rate
0.0
0.2
0.4
0.6
0.8
1.0
0
0.2
0.4
0.6
0.8
1

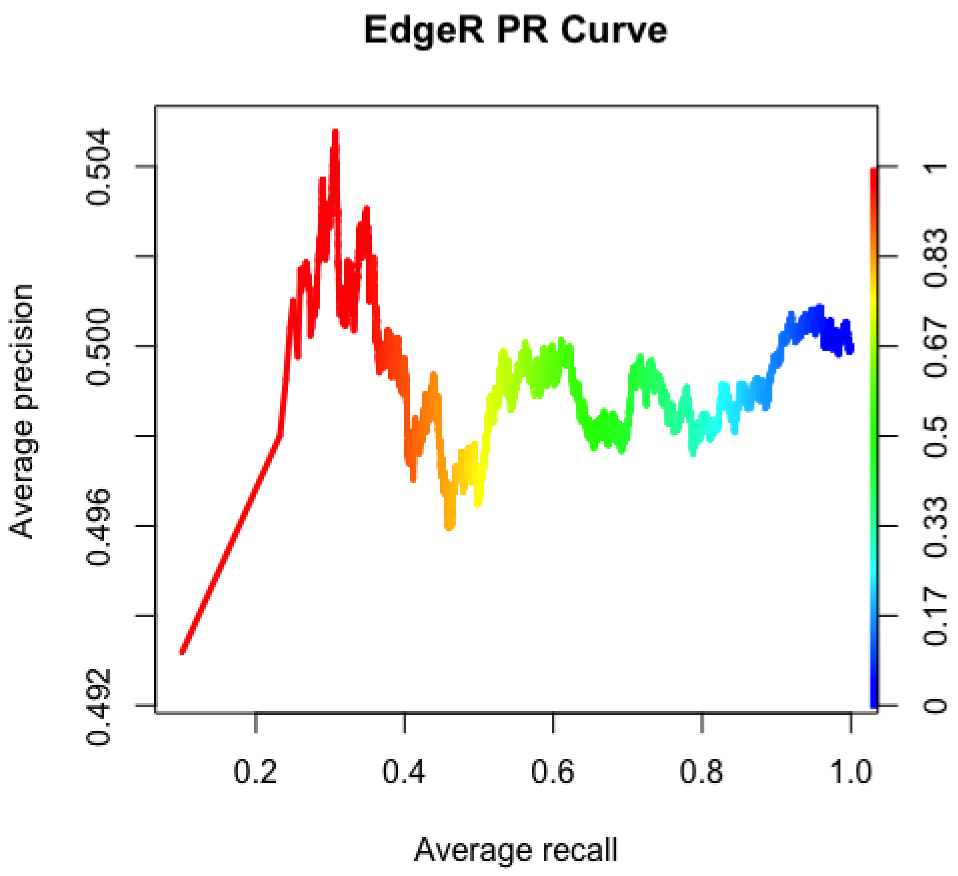


# QC

Check performance

```
## Confusion Matrix and Statistics
##
##           Reference
## Prediction    0    1
##          0 5962 3375
##          1 5952 3384
##
##                Accuracy : 0.5005
##                  95% CI : (0.4933, 0.5077)
##     No Information Rate : 0.638
##     P-Value [Acc > NIR] : 1
##
##                   Kappa : 0.001
##
##  Mcnemar's Test P-Value : <2e-16
##
##             Sensitivity : 0.5007
##             Specificity : 0.5004
##          Pos Pred Value : 0.3625
##          Neg Pred Value : 0.6385
##              Prevalence : 0.3620
##          Detection Rate : 0.1812
##    Detection Prevalence : 0.5000
##       Balanced Accuracy : 0.5005
##
##        'Positive' Class : 1
##
```

### make limma-voom QQ

check calibration

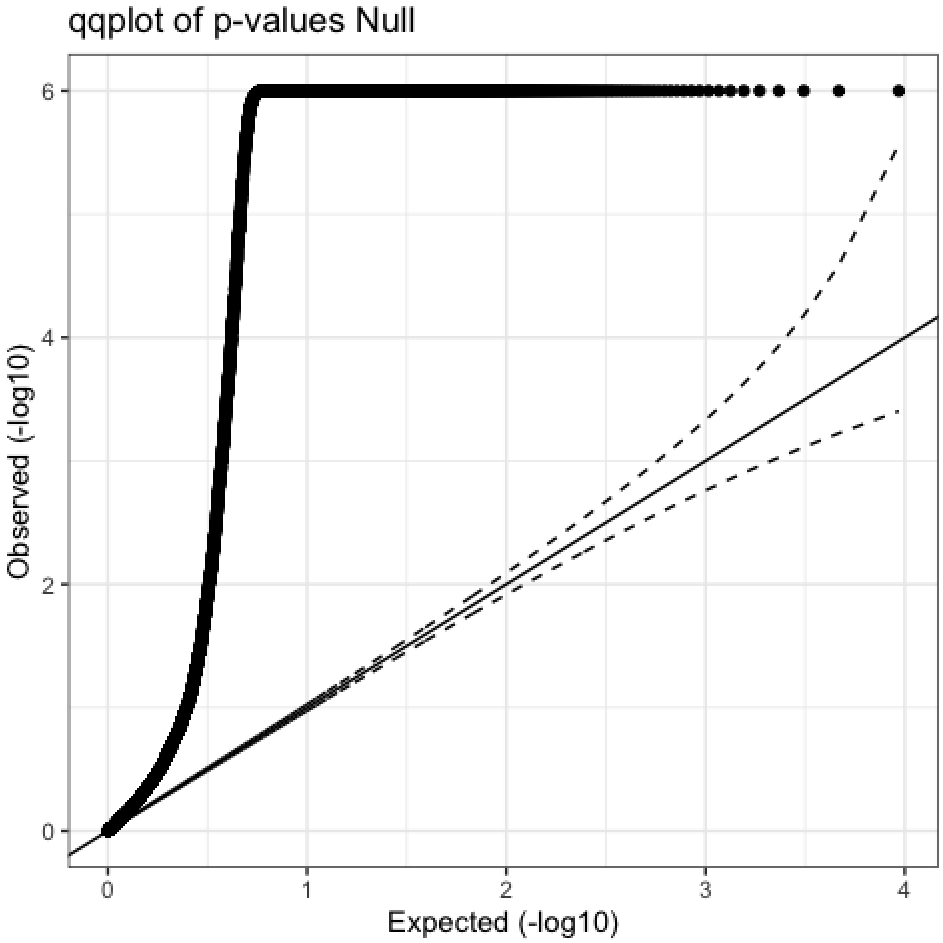


### run DESeq

Now run the differential expression test using DESeq2

```
## [1] "Intercept"   "cond_1_vs_0"
```

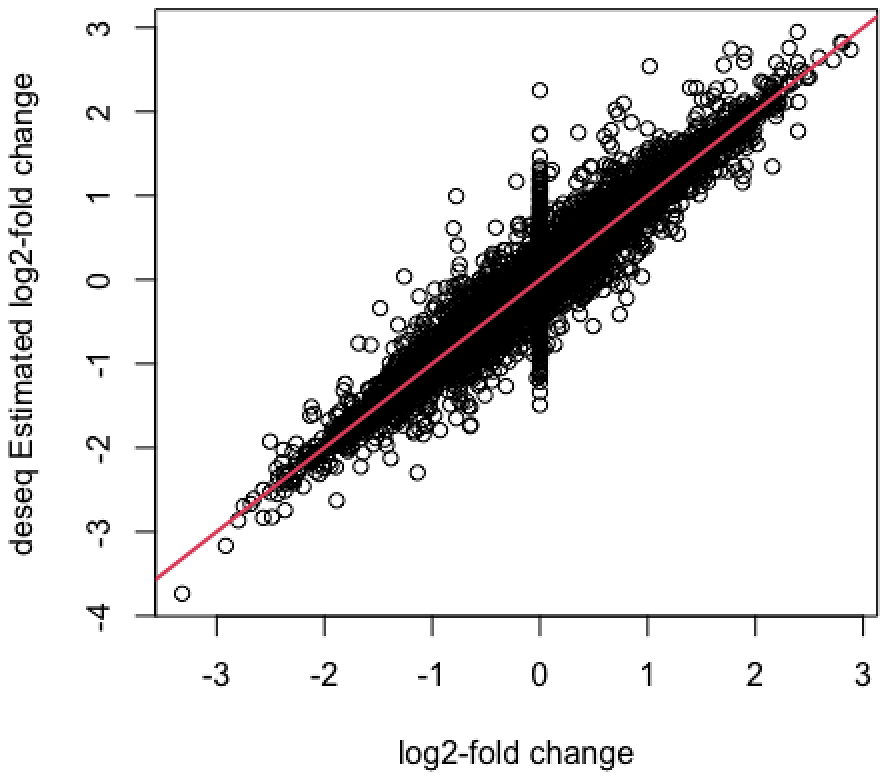
deseq Estimated log2-fold change
log2-fold change
-4
-3
-2
-1
0
1
2
3
-3
-2
-1
0
1
2
3

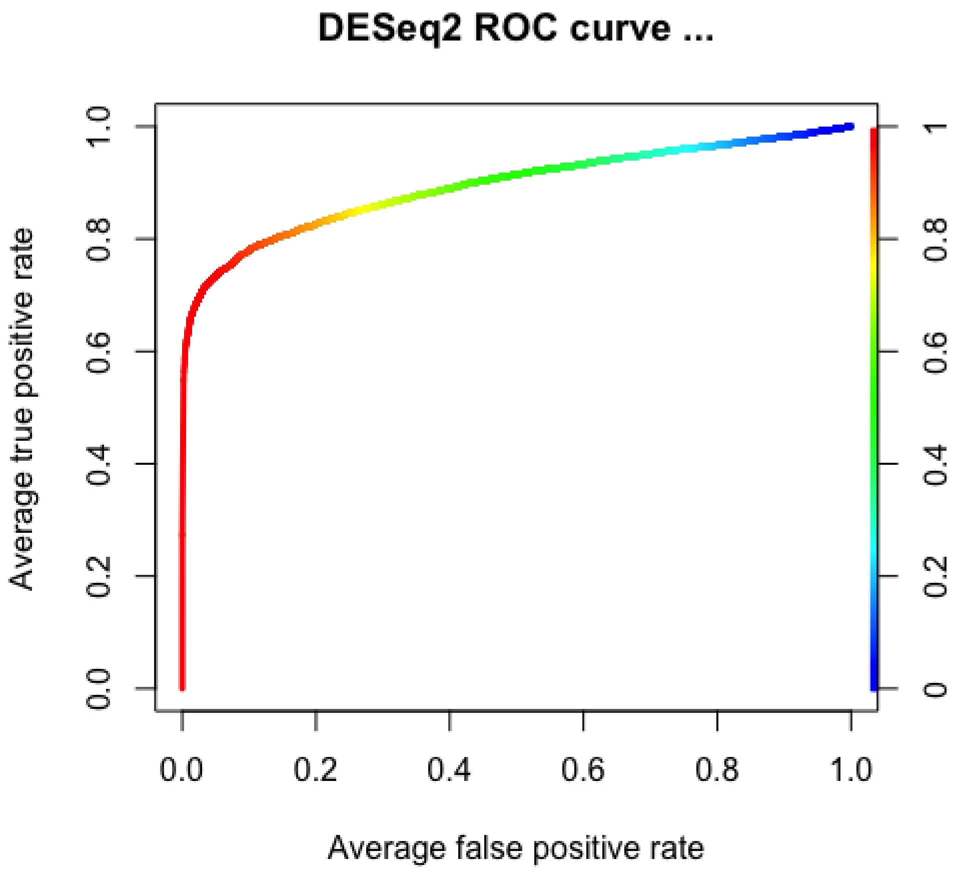
DESeq2 ROC curve ...
Average true positive rate
Average false positive rate
0.0
0.2
0.4
0.6
0.8
1.0
0.0
0.2
0.4
0.6
0.8
1.0
0
0.2
0.4
0.6
0.8
1

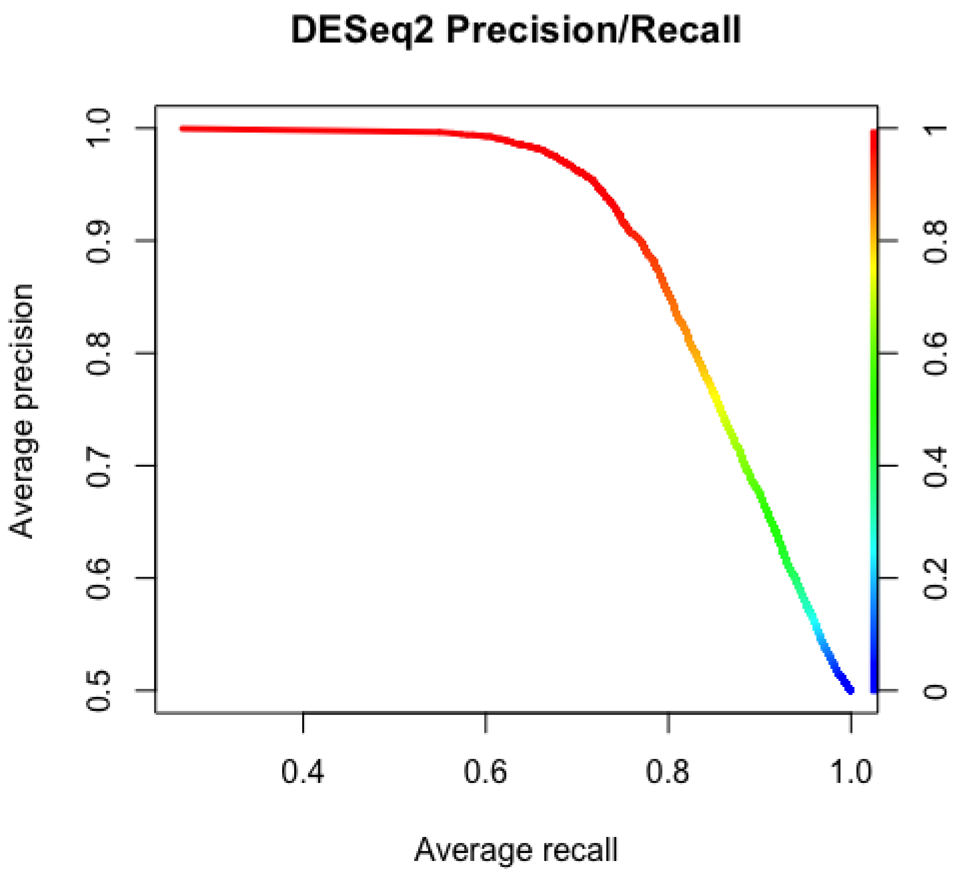


### QC DESeq

get performance

```
## Confusion Matrix and Statistics
##
##           Reference
## Prediction    0    1
##          0 8662  675
##          1 2297 7039
##
##                Accuracy : 0.8408
##                  95% CI : (0.8355, 0.8461)
##     No Information Rate : 0.5869
##     P-Value [Acc > NIR] : < 2.2e-16
##
##                   Kappa : 0.6817
##
##  Mcnemar's Test P-Value : < 2.2e-16
##
##             Sensitivity : 0.9125
##             Specificity : 0.7904
##          Pos Pred Value : 0.7540
##          Neg Pred Value : 0.9277
##              Prevalence : 0.4131
##          Detection Rate : 0.3770
##    Detection Prevalence : 0.5000
##       Balanced Accuracy : 0.8514
##
##        'Positive' Class : 1
##
```

### make DESeq2 QQ

check calibrations

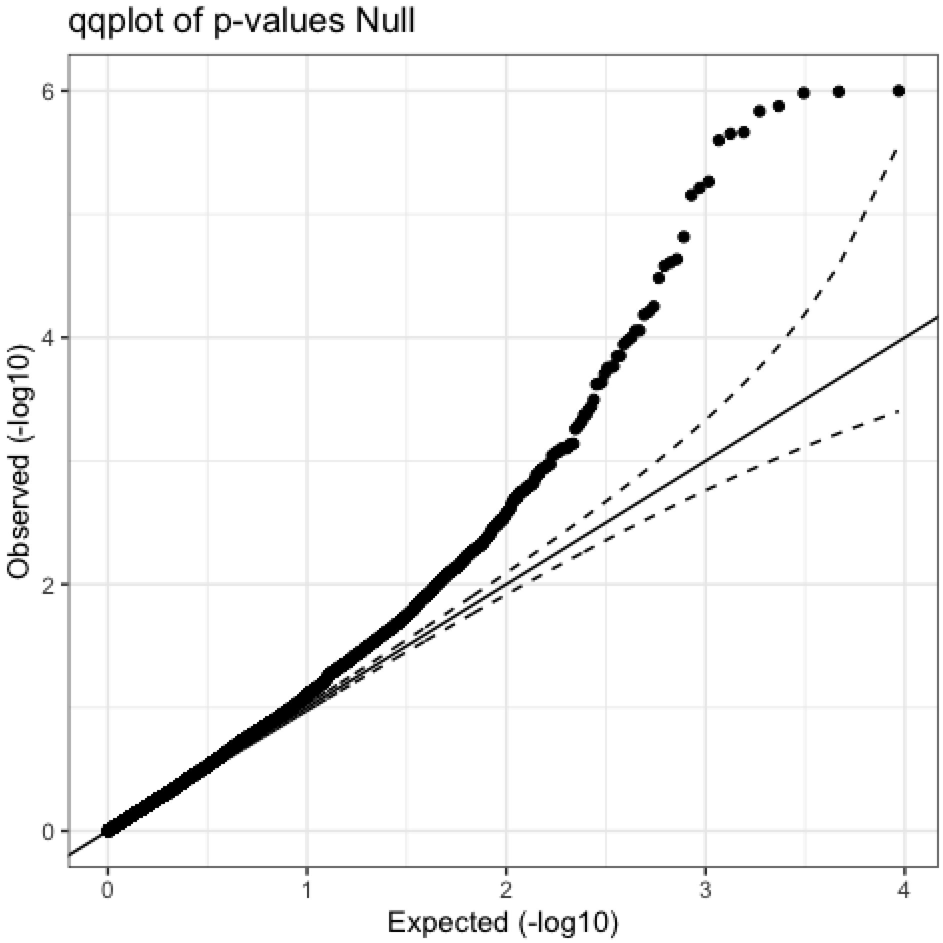


### Conclusion

In conclusion, we find that SAM performs equally as well as DESeq2 and outperforms edgeR. Therefore, we will keep using SAM and make sure we log10 the RNA-Seq counts.
